# How artists experience their own art

**DOI:** 10.64898/2026.03.31.715480

**Authors:** Giulia Tomasetig, Lucia Maria Sacheli, Margherita Adelaide Musco, Stefano Pizzi, Gianpaolo Basso, Grazia Fernanda Spitoni, Gabriella Bottini, Luigi Pizzamiglio, Eraldo Paulesu

**Author notes:** **Corresponding author**. (EP), (GT). These authors contributed equally to this work (Giulia Tomasetig, Lucia Maria Sacheli).

## Abstract

Humanity has always admired and created artwork, but the neurocognitive mechanisms behind artistic experience are still elusive. Professional artists and their intimate relationship with their artworks provide a unique opportunity to study the nature of art experience due to their expertise in both art making and art appreciation. During two fMRI tasks, professional artists (N=20) made aesthetic judgments on their own and other artists’ paintings (aesthetic appreciation task); they also mentally reconstructed the moments when they conceived their artworks or, as a control condition, when they visited now-familiar places for the first time (reconstruction by imagery task). During art appreciation of their own (as compared to other artists’) paintings, participants showed stronger recruitment of bilateral posterior parietal cortices, the left lateral occipitotemporal cortex, and the dorso-central sector of the right insula, that is, action-related brain regions also involved in encoding the emotional components of movements. The reconstruction of their own artistic creation (as compared to episodic memory retrieval) involved the left fronto-parietal network associated with motor cognition. Altogether, these results suggest that the mental representations of the actions involved in creating art are integral to the overall artistic experience of painters, supporting an embodied view of the artists’ experience of art.

## Introduction

From the earliest traces of human activity and across different cultures, humans have produced visual art (1). Artworks are universally admired and gathered by single individuals or social groups in private or public collections like museums or art galleries. Conceiving, realizing, or admiring artwork involves both cognitive processes and emotional engagement. As can be traced back in Nietzsche’s essay *The Birth of Tragedy,* the development of art is bound to the dichotomy of the Apollonian and the Dionysian (2). While Apollonian art emphasizes rationality and knowledge while curbing and repressing human impulses, Dionysian art relies on the corporal dimension and immediately and profoundly engages the senses, eluding rationality and conceptual learning (2).

Echoes of this dichotomy can also be found in the cognitive neuroscience of aesthetic perception and creative production. A more “Dionysian” framework suggests that aesthetic and creative experiences rely on corporal experiences and the automatic and pre-reflexive mechanism of motor and empathic resonance with the pieces of art or with the feelings and actions of the artist who created them (i.e., *Embodied Hypothesis*) (3). On the other hand, following a more “Apollonian” perspective, art appreciation can be seen as inherently linked to learning and knowledge acquisition. According to this view, aesthetic experience and creative expression require inhibiting ongoing physiological states, actions, and habitual thoughts, enabling new perceptual and conceptual insights to arise (i.e., *Perceptual Learning and Inhibitory Hypothesis*) (4).

More specifically, within the perceptual domain, the ***Embodied Hypothesis*** posits that the aesthetic experience of visual art relies on implicit motor simulation processes. These include the simulation of actions, emotions, and corporal sensations depicted in the works of art, as well as the gestures the artists performed in realizing the graphic marks on the canvas (3). The traces left by brushstrokes represent the visible outcome of motor acts: just as hearing sounds or viewing handwritten letters activates corresponding motor programs in the listener or viewer (5–8), the observation of brushstrokes is thought to evoke in the beholder a sense of bodily resonance with the movements implied by those physical traces (3). Similarly, bodily sensations have been proposed as central to the aesthetic experience (9). The *Embodied Hypothesis* also highlights the role of motor simulation in artistic creativity and production. The simulation of possible future actions would support improvising and generating creative ideas (10).

Instead, a different theoretical framework, an ad-hoc declension of the Predictive Coding Hypothesis (11–13), de-emphasizes the role of motor simulation in aesthetic appreciation in favor of the processing of sensory stimuli that are not adequately explained by predictions and yet carry potential learnable information. According to some, this would go in parallel with the inhibition of motor behavior and a simultaneous attentional focus on the perceptual features of the stimuli (4). From this perspective, motor inhibition would allow prediction updating, thereby fostering learning dynamics and knowledge acquisition. Aesthetic pleasure would thus represent the result of a switch from a state of higher unpredictability – that captures our attention – to a state of higher predictability (i.e., uncertainty resolution), facilitated by inhibitory control in the first place (4). Concerning creativity, the ability to inhibit the most common and automatic but irrelevant or interfering responses, such as obvious or conventional associations, would contribute to the generation of innovative ideas for new creations (14,15). In what follows, this hypothesis will be called collectively the ***Perceptual Learning and Inhibitory Hypothesis*.**

Neuroscientific studies in the domains of visual aesthetics and creativity have mostly been exploratory and not guided by neurocognitive models (16). Moreover, all previous studies on aesthetic appreciation tried to bypass the effect of artistic education, including only participants naïve to the history or practice of visual art and presenting little-known paintings (17). The study of professional artists, instead, provides a unique opportunity to investigate within the same participants the neurocognitive bases of both art-making and art-viewing, thus providing a more comprehensive testbed on the psychophysiology of art experience. While a recent meta-analysis of fMRI studies on aesthetic appreciation and creativity in naïve onlookers seems to support the *Perceptual Learning and Inhibitory Hypothesis* mentioned above (17), the study of professional artists may represent a perfect test-case to assess whether this hypothesis can be generalized or whether professional artists experience art in a more embodied manner, independently from the subject and its figurative potential of evoking motor acts, because of their *unique artistic motor repertoire*, as discussed below.

### Aims of the study and neurofunctional predictions

The present fMRI study aimed to investigate the neurocognitive processes underlying professional artists’ aesthetic and creative experiences in the visual domain, while also challenging the *Embodied* and the *Perceptual Learning and Inhibitory Hypotheses*. For this purpose, we recruited expert painters who performed two different fMRI tasks. In one task, they observed and mentally expressed aesthetic judgments on their own paintings and paintings made by other artists (i.e., aesthetic appreciation task). In the other task, they had to reconstruct what they thought and felt when they conceived some of their own pieces of art or, as a control task, when they visited now familiar places for the first time (i.e., reconstruction by imagery task). Importantly, the selection of places, own and others’ artworks was individualized for each artist and fully matched for trivial differences (e.g., artistic style; see the Preliminary Phase below). We planned to match aesthetic judgments across the different classes of stimuli shown during fMRI (i.e., 20 own paintings, 20 others’ paintings, and 20 places). To this aim, aesthetic judgments were collected in a first behavioral session (Day 1, see below). This procedure ensured that any differences observed in neural responses to one’s own vs. others’ paintings or between the reconstruction by imagery of the creation of one’s own artworks vs. the visit to familiar places could not be attributed to variations in subjective liking. Own and others’ artworks were also matched for style, content, and dynamism. We made sure that artists were highly familiar also with paintings made by others.

Comparing artists’ responses to their own versus others’ paintings allowed us to investigate the role of embodied vs. predictive and inhibitory processes, as the two theoretical standpoints lead to opposite predictions regarding how this factor should modulate the artists’ neurofunctional responses. On the one hand, people can recognize and discriminate the products of their past actions from those generated by others (18,19). This so-called “authorship effect” may be driven by the perception of self-kinematic motor information embedded in static traces, rather than by visual familiarity alone (20,21) or episodic memory (22). The hypothesis was that the motor system may resonate more strongly with signals stemming from prior self-produced actions (19,23). Thus, self-generated paintings should elicit motor representations of gestures previously enacted by artists during their creation.

From the *Perceptual Learning and Inhibitory Hypothesis* perspective, individuals can generate better predictions for forthcoming visual and auditory outcomes for their own actions compared with those of others (18,23). Coherently, the outcomes of self-generated actions should be usually associated with sensory attenuation – reported both at the phenomenological and neural level in primary sensory brain regions – an event commonly attributed to predictive processes (24,25), while observing others’ paintings should reflect in a major perceptual uncertainty, leading to stronger signals in unimodal visual brain regions with a concurrent action inhibition to enhance perceptual learning.

Thus, according to the *Embodied Hypothesis,* professional artists should show stronger recruitment of brain regions involved in motor resonance while observing their own vs. others’ paintings, as well as while reconstructing creative moments, compared with the control condition. Instead, the *Perceptual Learning and Inhibitory Hypothesis* predicts greater recruitment of perceptual and motor inhibitory brain regions when viewing others’ than one’s own paintings, as well as the engagement of brain regions involved in inhibitory control during the reconstruction of the creative moments.

## Materials and methods

### Participants

Twenty-eight professional artists participated in the study’s first behavioral session (Day 1, see below). However, four of them did not join in the second fMRI session (Day 2) and other four were excluded due to areas of altered signal intensity revealed by the MRI, thus leaving a final group of twenty participants (10 females, 14 right-handed, 4 ambidextrous, age range 22-74 years, mean age 42.15 ± 18.81), all with high education (range 16-18 years). The sample size is consistent with previous fMRI studies on aesthetic appreciation and creativity: across the thirty-four experiments included in a recent meta-analysis on the same topics, the median sample size is twenty participants (17) All the participants had normal or corrected to normal vision, and none reported a history of neurological or psychiatric disorder. All the artists were screened to exclude age-related neurocognitive decay by using the Mini-Mental State Examination (MMSE) (26); no one showed pathological performance after correcting the raw scores for age and education (MMSE corrected score range 25.85 – 30). The experimental protocol was approved by the Ethics Committee of the University of Milano-Bicocca, where data collection took place (from May 2, 2022 to July 19, 2023), and the study was conducted in accordance with the Declaration of Helsinki. Written informed consent was obtained from all participants.

Inevitably, our experiment did not rely on comparisons with a control group for several principled reasons:

(1) The core task in this study involves artists viewing and reflecting on artworks they created themselves. This kind of self-referential engagement requires authorship, which, by definition, non-artists lack, together with a broad body of artworks usable as stimuli.
(2) Asking non-artists to reflect on “their own artworks” would require fabricated or trivial creations, which would lack the motor, emotional, and aesthetic depth central to the phenomenon under study.
(3) Not all questions require control groups: control groups are essential when isolating variables in general cognitive mechanisms, but this study addresses a specific expert population and experience. The aim was not to determine what all humans do, but to uncover how expertise and authorship uniquely shape neurocognitive responses in experienced artists.

### Procedure

The study was divided into three phases: a preliminary phase, a first behavioral session (Day 1), and a second fMRI session (Day 2).

#### Preliminary phase

Participants were professional artists selected by an artist and professor of the Academy of Fine Arts of Brera (the author S.P.). They were recruited via e-mail or phone call, during which they were informed about the purposes of the study. If they consented to participate, they were asked to send forty images of their own pictorial artworks and forty pictures of places they knew. For each participant, S.P. provided a list of renowned artists with the same pictorial style, and two independent experimenters (the authors G.T. and M.A.M.) selected a “control” painting for each artwork (i.e., a one-to-one matching with the forty self-selected works of art by each artist) according to the following criteria: i) abstract/figurative; ii) living/non-living; iii) static/dynamic; iv) canvas coverage (very/little covered); v) very/little colored; vi) pictorial technique. For each artwork of each participant, a third independent researcher (the author L.M.S.) selected the best “control” painting out of the two options.

#### Day 1

In the first session, the artists performed two behavioral judgment tasks. In the first one, they saw their own forty and other artists’ forty paintings and had to provide liking judgments on a Visual Analogue Scale (VAS) ranging from 0 to 100. In the second task, they evaluated the grade of appreciation of the forty places using the same VAS (see S1 Fig in the Supporting Information for the trial-timeline). For both tasks, each image was presented twice to obtain a more robust evaluation. During the same session, the artists were also screened for contraindications to fMRI. For each subject and class of stimuli (i.e., own paintings, others’ paintings, and places), we calculated the mean score from the two evaluations of each image. We then selected twenty items for each category by including the ten least and ten most liked items, while also ensuring that there were no significant differences in ratings across classes of stimuli (own mean: 63.68 ± 24.53; others mean: 63.69 ± 24.57; places mean: 64 ± 26.43). The twofold aim of this procedure was (i) to ensure that any differences across classes of stimuli were not due to differences in liking, and (ii) to evaluate in a further analysis whether the artists’ brain reacts differently to paintings that they like the most compared to those they liked the least. For each artist, their own and other artists’ selected artworks were also matched for aspects related to the content (i.e., number of figurative and abstract artworks, number of paintings carrying body-related cues) and the dynamism (i.e., number of static and dynamic artworks). The sixty selected images (twenty own paintings, twenty other artists’ paintings, and twenty places) were used in two fMRI tasks.

#### Day 2

In the second session, participants completed two fMRI tasks in two different runs: an aesthetic appreciation task and a reconstruction by imagery task. The order of the tasks was counterbalanced between participants. Prior to the scanning session, the artists completed a brief training using images that were not selected to familiarize themselves with the tasks.

During the **aesthetic appreciation task**, participants observed both their own and others’ artworks and were instructed to mentally formulate an aesthetic judgment. No motor response was required to avoid confounding motor-related activations associated with response preparation and execution, which could, in principle, amplify motor signals and artificially bias the data toward an embodied interpretation. However, to ensure participants remained focused and engaged in the task, and keep sufficient control on their performance, a question randomly appeared on the screen, prompting them to press a button with either their index finger or thumb depending on whether the artwork they had just seen belonged to another artist (index finger) or if it was their own (thumb; see Fig 1a for the trial timeline).

**Fig 1.**
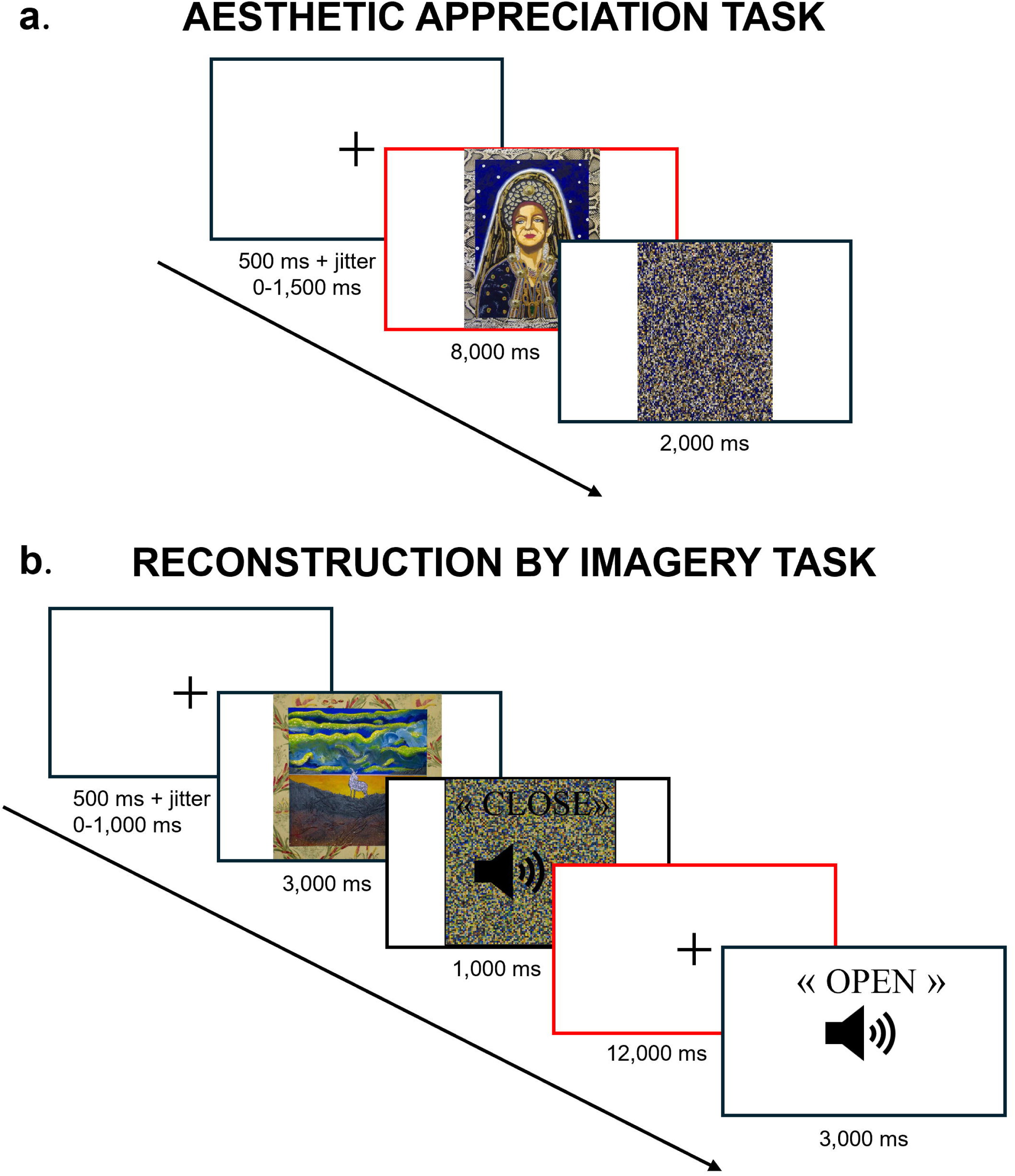
Trial-timeline of the experimental tasks. The figure illustrates the trial-time line of the a) aesthetic appreciation task, and the b) reconstruction by imagery task. The red frames indicate the onset of the event of interest in the fMRI analyses. For illustrative purposes, we report two paintings of the artist Stefano Pizzi.

During the **reconstruction by imagery task,** artists were instructed to close their eyes and mentally evoke:(i) the moment when they conceived their own artworks, when one of their paintings appeared on the screen, or (ii) the moment when they visited the places for the first time, when an image of a place appeared (control condition). They were asked to press a button either with the index or thumb finger when the recall was vivid (see Fig 1b for the trial-timeline). We designed the control condition in order to isolate the neurofunctional underpinnings of artistic production from those more generally involved in episodic memory retrieval.

Once outside the scanner, we asked the artists to qualitatively describe how they performed the aesthetic appreciation and reconstruction by imagery tasks. Then, i) they rated the imaginability of each own artwork and place (i.e., how well they imagined and evoked the moments during the reconstruction by imagery task) on a VAS, and ii) they rated their emotional involvement with each stimulus (own and others’ paintings and places) also using a VAS.

### Stimuli and trial timeline

During the fMRI session, the visual and auditory stimuli were delivered through MRI-compatible devices (NordicNeuroLab InRoom Viewing Device and Audio System). The aspect ratio of the visual stimuli was preserved by adjusting the image size to a maximum dimension of 1920 × 1080 pixels. The responses were recorded through an MRI-compatible response-box (NordicNeurolab ResponseGrip) placed under the right hand of the participants.

The **aesthetic appreciation task** comprised ten mini-blocks of six trials each (for a total duration of ∼ 10 min). Each mini-block comprised four paintings (i.e., two of their own paintings and two of others), a question (4,000 ms), and an implicit baseline (i.e., black screen displayed for 12,000 ms) presented in random order. The trials started with a fixation-cross displayed on the screen (variable duration ranging from 500 to 2,000 ms). The image of an artwork was then presented (8,000 ms), followed by its scrambled version (2,000 ms, see Fig 1a).

The **reconstruction by imagery task** comprised ten mini-blocks of five trials each (for a total duration of ∼ 14 min). Each mini-block comprised four images (i.e., two of their own paintings and two places) and an implicit baseline (i.e., black screen displayed for 12,000 ms) presented in random order. Each trial started with a fixation-cross (duration ranging from 500 to 1,500 ms), followed by the image of one of their paintings or a place (3,000 ms). Then, simultaneously, the scrambled version of the image appeared, and participants heard “close” (1,000 ms). After they had closed their eyes, the participants had a maximum of 12,000 ms to evoke what appeared in their mind and what they felt when they conceived their own artworks or when they visited places for the first time. If they took a shorter time, they were instructed to press a button with their index or thumb finger when the recall was vivid. Anyway, they had to remain with their eyes closed for the entire duration (12,000 ms), until they heard “open” (1,000 ms), i.e., the instruction to open their eyes and interrupt the mental reconstruction (see Fig1b). This procedure allowed us to collect the same number of scans across participants, although participants took different durations to complete the trials. The software E-Prime 3 (Psychology Software Tools, Inc.) was used both for stimuli presentation and randomization.

### Behavioral Analyses

All behavioral data are available in the Open Science Framework at the following address: https://osf.io/e4fdb/.

#### Day 1

Based on the liking judgments provided by the artists on Day 1, we selected the stimuli for the fMRI experiments (i.e., artists’ own paintings, other artists’ paintings, and places). The stimuli were matched for liking ratings across categories, as ensured by the results of a three-way (own artworks/others’ artworks/places) within-subject non-parametric ANOVA (i.e., Friedman test) performed for each artist (all *p* values were greater than 0.063). Additionally, for each artist, we checked that aspects related to content and dynamism were matched between the artists’ own and others’ artworks. We indeed classified each artwork based on whether i) its content was figurative or abstract, ii) it carried body-related cues or not, and ii) its configuration was dynamic or static, and we checked that the frequencies of these features did not differ significantly between own and others’ paintings, as ensured by the results of a Chi-Squared test performed for each artist (all *p* values were greater than 0.058).

#### Day 2

At the behavioral level, we wanted to check that there were no significant differences in the duration of the mental reconstruction during the fMRI task, nor in imaginability between own paintings and places to avoid entering confounds in the analysis that could explain some of the differences in the brain activations. From the behavioral analysis on duration of mental reconstruction, we excluded one subject because, although she attempted to press the button, no response was recorded from the response box. For this participant, duration was set equal to 0 for the fMRI analyses in all trials. Then, we explored whether there were differences in emotional involvement across the three classes of stimuli. All these analyses were performed using linear mixed effects models with stimuli as fixed effects. By-subjects random intercepts and slopes were included to account for variability across subjects. Data were analyzed using the statistical software Jamovi (retrieved from https://www.jamovi.org.) and the GAMLj module (retrieved from https://gamlj.github.io/). Degrees of freedom were estimated using the Satterthwaite method. Post hoc test applied the Bonferroni correction when needed, and all inferential tests of significance were based upon an α level of .05.

### MRI data acquisition and analyses

#### MRI data acquisition

Whole-brain functional T2*-weighted images were acquired using a 3.0-T Ingenia CX scanner (Philips), equipped with gradient-echo echo-planar imaging. The T2*-weighted brain volumes were acquired with the following parameters: repetition time [TR] 2000 ms, echo time [TE] 30 ms. Each volume comprised 35 transversal slices, acquired in a descending, not-interleaved order; each brain slice was 4 mm thick with no interslice gap. The flip angle [FA] was 75° with a field of view [FOV] of 240 mm, matrix size 80 × 80. Overall, 305 scans were acquired for the aesthetic appreciation task and 435 scans for the reconstruction by imagery task. A high-resolution T1-weighted structural image (TR 8.1 ms, TE 3.7 ms, 207 sagittal slices, FOV 256 mm, matrix size 256 × 256, FA = 8°, Inversion Time 1000 ms) was also acquired for each participant.

#### Preprocessing

After image reconstruction, we converted raw data from the DICOM to the NIfTI format using the MRIconvert software. All subsequent data analyses were performed in MATLAB R2019b (MathWorks) using Statistical Parametric Mapping software (SPM12, Wellcome Centre for Human Neuroimaging, UCL). For each participant, we first spatially realigned and unwarped fMRI scans to the first volume of each run to correct for head movement artefact. The T1-weighted structural image of each participant was segmented and warped into the normalized space (SPM12 template, tmp.nii); it was used as a reference for the coregistration with the unwarped images, and the deformation fields saved from T1 segmentation were then applied to the coregistered functional scans. The data matrix was interpolated to produce voxels with 2 × 2 × 2 mm dimensions. The normalized scans were finally smoothed using a Gaussian filter of 10 × 10 × 10 mm to improve the signal-to-noise ratio and allow the application of the family-wise error rate (FWER) cluster-level correction in the group-level univariate analyses (27).

We also used the Artifact Detection Tools (ART, http://www.nitrc.org/projects/artifact_detect) to identify outlier scans for global signal and movement. We marked time-points as outliers if scan-to-scan variations in the global signal exceeded 3SD from the mean, and if the compounded measure of movement parameters exceeded 1 mm scan-to-scan movement average. We accounted for these outlier datapoints by adding specific regressors of no interest at the stage of the single-subject analysis. These outlier volumes were 2.72% ± 2.46% in the aesthetic appreciation task and 4.32% ± 3.98% in the reconstruction by imagery task. No participant was excluded from the analyses due to excessive movement, as established by the threshold of 20% of outlier scans for each experimental condition in the fMRI run.

#### Inferential strategy and statistical analyses of the fMRI data

The inferential strategy adopted in this study was straightforward: first, to assess whether a null hypothesis – namely, that no difference was present between experimental and control stimuli – could be rejected. In principle, this was anything but trivial, particularly for the aesthetic appreciation task, given that the stimuli – own and others’ artworks – were matched as tightly as possible. If the null hypothesis was rejected, the data would then be scrutinized using quantitative reverse inference as permitted by the software/database Neurosynth, which permits quantifying the functional association of a given brain map with functional/psychological terms. Neurosynth is widely used and, in our case, could have served well to decide which functional hypothesis of art appreciation – the *Embodied Hypothesis* or the *Perceptual Learning and Inhibitory Hypothesis* – was more likely. We also reasoned that a congruent result with the reconstruction by imagery of art creation would corroborate the interpretation of the art appreciation data (see below).

The statistical analyses were based on canonical univariate analyses because they are fully transparent and because the structure of the events was such that any multivariate classification attempt was unfeasible given the length of each event and their numerosity.

A two-step statistical analysis based on the general linear model (GLM) was performed. At the first level of analyses (single-subject level), the blood oxygen level-dependent (BOLD) signal associated with each experimental condition was analyzed by convolution with a canonical hemodynamic response function (28). Global differences between the fMRI time-series were removed from all voxels with grand mean scaling (global normalization). The time series was high-pass filtered at 128 s and pre-whitened by means of an autoregressive model AR(1) to remove artifactual contributions to the fMRI signal, such as noise from cardiac and respiratory cycles. This first step implied a fixed-effect analysis, in which condition-specific effects were calculated in each participant. We separately analyzed the aesthetic appreciation and reconstruction by imagery runs, as reported below.

In the **aesthetic appreciation** run, we characterized the BOLD signal associated with observation of own and others’ paintings in each trial (event-related design). We thus modeled two regressors of interest (others and own). The event duration was set to 8 s, and the onset of the event at each trial corresponded to the onset of the image. Separate regressors modeled experimental confounds, including i) trials in which participants responded to the questions, ii) the signal associated with the observation of the scrambled images, iii) the realigning parameters calculated in the preprocessing step, and iv) a final separate regressor for each outlier scan to be removed from the analysis.

Concerning the **reconstruction by imagery** run, we modeled two regressors of interest (places and own paintings) in an event-related design. The event duration was calculated as the time-window between the instructions to start the reconstruction task (i.e., black screen) after having closed their eyes and the instant when the artists pressed the button to indicate the task was completed (we subtracted 2 s from this instant to avoid activations related to movement preparation). When the participants did not press one of the two buttons, the duration was set to 11 s. The onset of the event at each trial corresponded to the onset of the black screen that followed the instruction to close their eyes and start the reconstruction task. As confounds, we modeled in separate regressors the signal associated with i) the observation of the photograph depicting paintings or places and the listening of the instruction to close the eyes, ii) the listening of the instruction to open the eyes, and iii) the execution of a motor response. The realigning parameters were also entered as regressors of noninterest, as well as a final separate regressor for each outlier scan to be removed from the analysis.

After the design specification and estimation, we characterized the effects associated with each experimental condition, applying a weight of +1 for the regressor-of-interest and a weight of zero for all other regressors. The two effects related to aesthetic appreciation and the two related to reconstruction by imagery were entered into two separate second-level dependent sample t-tests that conformed to random effect analyses. One second-level analysis included painting (others vs own) as a within-subjects factor, while the other included stimuli (places vs own paintings) as a within-subjects factor. Concerning the analyses on aesthetic appreciation, to ensure finding clusters of real activations (i.e., not driven by deactivations in the opposite condition), we masked the contrast “own > others” with the simple effect of activation related to the observation of own paintings (“own > implicit baseline”) and the contrast “others > own” with the simple effect of activation related to the observation of others’ paintings (“others > implicit baseline”). The mask threshold was set at *p*_uncorr_ < 0.05 at the voxel-level. In the same way, we masked the contrast “own paintings > places” with the simple effect of activation related to the reconstruction of own paintings’ creation (“own paintings > implicit baseline”, *p*_uncorr_ < 0.05 at the voxel-level) in the analysis on reconstruction by imagery.

We conducted a further fMRI analysis to examine whether liking had an effect on aesthetic appreciation. We separately analyzed the own and others’ artworks that obtained the best and the worst judgments on Day 1 (own less-liked mean: 43.92 ± 18.14; others’ less-liked mean: 43.95 ± 18.23; own liked mean: 83.43 ± 9.63; others’ liked mean: 83.42 ± 9.75). Thus, we entered liking (less-liked/liked) as a further within-subjects factor in the fMRI analysis. We modeled four regressors of interest and the corresponding effects at the first level of analysis (own less-liked, own liked, others less-liked, others liked). The four effects were entered into a second-level full-factorial analysis of variance (ANOVA) that included paintings (own vs others) and liking (less-liked vs liked) as within-subject factors. Analyses were conducted at the whole-brain level. The voxel-wise threshold applied to the statistical maps before the cluster-wise correction was *p_uncorr_* < 0.001, as recommended by Flandin and Friston (27). We then report in the tables in the Main Text: (i) all peaks included in the clusters that survived family-wise error rate (FWER) correction for multiple comparison at the cluster-level, (ii) all peaks that survived a FWER-correction at the voxel level, and (iii) for the sake of clarity, the peaks included in cluster significant at *p*_uncorr_ < 0.05 (but the application of a lower threshold in these instances is highlighted in the captions).

All neurofunctional data are available in the Open Science Framework at the following address: https://osf.io/e4fdb/.

### Neurosynth decoding

To further decipher the functional role of the brain regions resulting from our analyses, we used the decoder function implemented on the website Neurosynth (https://neurosynth.org, (29). It permits quantifying the functional association of a given brain map with functional/psychological terms, after interrogation of 14,371 functional MRI studies and their semantics. We conducted two decoding analyses: in one, we entered as input image the brain map resulting from the direct comparison between the appreciation of one’s own and others’ artworks (own > others) masked with the simple effect related to the appreciation of one’s own paintings. In the other one, we entered as input image the brain map resulting from the reconstruction of art creation compared with the control task (own paintings > places) masked with the simple effect related to the reconstruction of own paintings’ creation.

## Results

### Behavioral results

Concerning the imaginability, we did not find significant differences between own paintings and places (F (1,19) = 1.02, *p* = 0.326; respectively, M = 76.2, SE = 3.05 and M = 78.7, SE = 2.23. See Supporting Information S2a Fig). Similarly, participants did not take different durations in recalling the creative moments compared to the ones when they visited the places (F(1,17.5) = 0.323, *p* = 0.577. Respectively, M = 9.68, SE = 0.387; M = 9.58, SE = 0.4. See Supporting Information S2b Fig). Finally, regarding emotional involvement, we found a significant main effect of the class of stimuli (F(2,19) = 26.8, *p* < 0.001). Post-hoc comparisons revealed that participants were less emotionally engaged with others’ paintings than with their own paintings and places (respectively, t(19) = −7.284, *p_corr_* < 0.001 and t(19) = −5.648, *p_corr_* < 0.001; M_others_ = 50, SE = 3.33; M_own_ = 70.5, SE = 2.28; M_places_ = 69.8, SE = 2.26). Instead, no differences emerged between own paintings and familiar places (t(19) = 0.284, *p_corr_* > 0.99. See Supporting Information S2c Fig).

### fMRI results

The simple effects (task versus implicit baseline) and the common effects in the aesthetic appreciation task and the reconstruction by imagery task are reported in the Supporting Information (see S3 Fig, S1-S3 Tables and S4 Fig, S4-S6 Tables respectively).

#### Aesthetic appreciation task

The direct comparison of the two appreciation conditions (for self-authored and for artworks made by others) revealed that in professional artists the appreciation of their own paintings was associated with stronger recruitment of the right superior parietal lobule (SPL) and anterior intraparietal sulcus (AIP), the left lateral occipitotemporal cortex (LOTc), and the dorso-central sector of the right insula (Fig 2a and Table 1). A left superior parietal cluster also showed a similar trend (k = 395, *p_uncorr_* = 0.019 cluster-level; local maximum at MNI −18, −60, 44; z-score = 3.8; *p_uncorr_* = 0.00007 voxel level).

**Fig 2.**
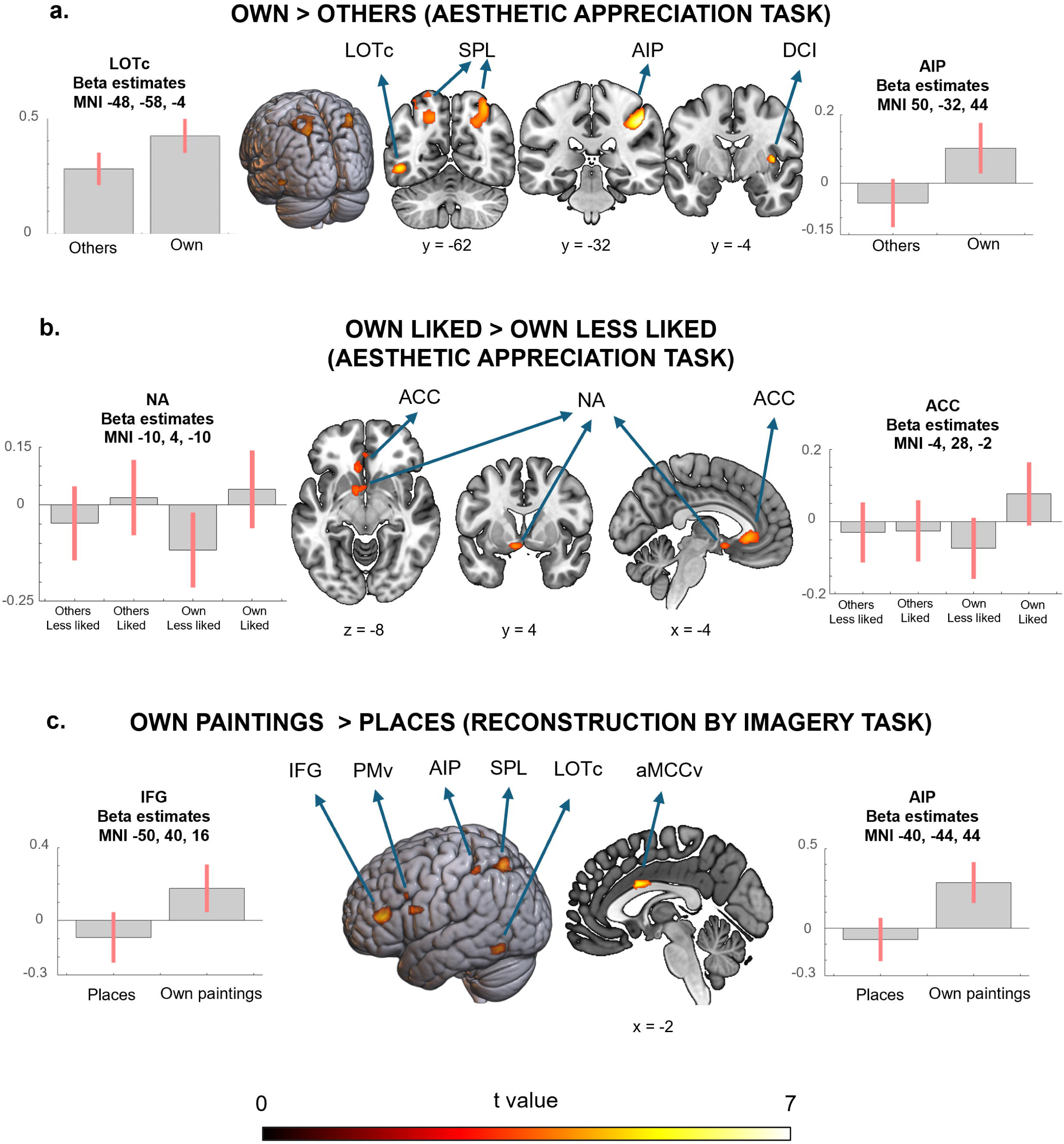
FMRI results. The figure illustrates a) the brain regions that show a greater engagement during the aesthetic appreciation of own compared with others’ artworks, b) the clusters that shows a greater engagement during the observation of own artworks that are more appreciated (Own Liked) compared with own artworks that are less appreciated (Own Less liked), and c) the cerebral regions that exhibit a greater involvement during the reconstruction of the creative moments compared with the control task. All the analyses were conducted at the whole-brain level, and the data are reported by applying the same statistical threshold discussed in the text (*p*_uncorr_ < .001 at the voxel level and *p*_FWER-corr_ < 0.05 at the cluster or voxel level). For illustrative purposes, we also report the left superior parietal cluster (own > others, aesthetic appreciation task), the anterior cingulate cortex, the nucleus accumbens (own liked > own less liked, aesthetic appreciation task), and the left ventral premotor cluster (own paintings > places, reconstruction by imagery task), though they do not survive the correction for multiple comparisons. LOTc = lateral occipito-temporal cortex; SPL = superior parietal lobule; AIP = anterior intraparietal sulcus; DCI = dorso-central insula; ACC = anterior cingulate cortex, NA = nucleus accumbens; IFG = inferior frontal gyrus; PMv = ventral premotor cortex; aMCCv = anterior-ventral midcingulate.

**Table 1.**
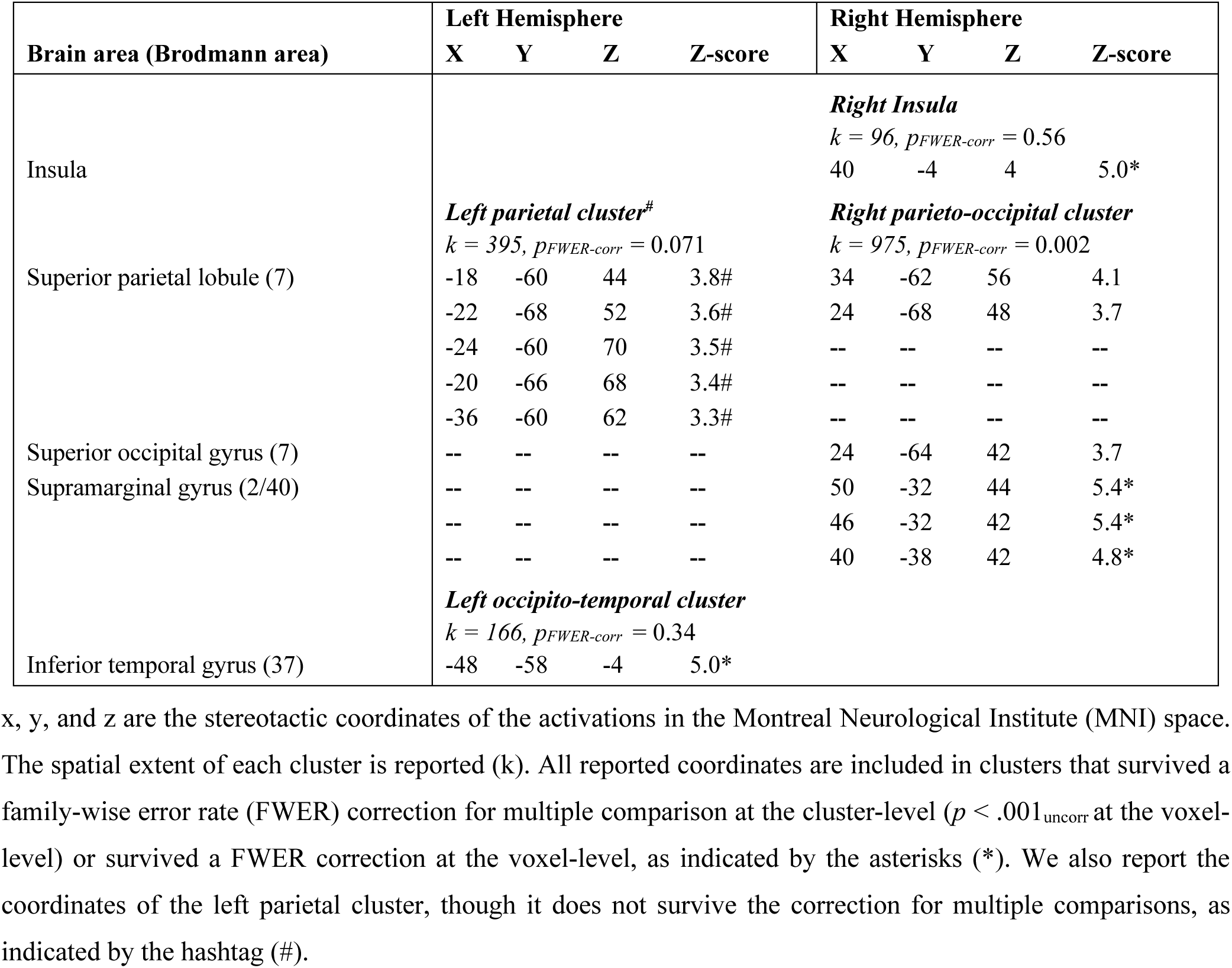
Neurofunctional results of the aesthetic appreciation task (“own > others” masked by the simple effect of own artworks).

Instead, the contrast in the opposite direction (“others > own”) did not show any significant effect.

We also assessed that the neurofunctional differences between the appreciation of their own paintings and paintings made by other artists were not driven by the behavioral differences in the emotional involvement, performing a control analysis with the delta of the emotional involvement (own – others) as a covariate of a one-sample t-test in which we entered the contrast “own > others” calculated at the first level of analysis. These results confirmed that differences in neural activations were not due to differences in emotional engagement (see Supporting Information S7 Table).

Moreover, we were interested in evaluating whether the brains of our artists reacted differently to artworks they liked more. For the simple effects of “liked > less liked” for own or others’ paintings, we found a cluster comprising the ventral anterior cingulate cortex (vACC) and the left nucleus accumbens only for liked own paintings (vACC, k = 306, *p*_uncorr_ = 0.044 cluster-level. Local maxima: MNI −4, 28, −2; z-score = 4.28; *p*_uncorr_ = 0.000000009 at the voxel-level, and MNI 0, 38, −8; z-score = 3.39; *p*_uncorr_ = 0.000347 voxel-level. Left nucleus accumbens, k = 95, *p*_uncorr_ = 0.239 cluster-level. Local maximum: MNI −10, 4, −10; z-score = 3.92; *p_uncorr_* = 0.00004 voxel-level) (see Fig 2b). On the contrary, not such liking-related functional anatomical effect was present during the observation of artworks made by someone else, whether they liked them or not.

#### Reconstruction by imagery task

Mental reconstruction of the creative moments compared with the control task was associated with greater recruitment of the left SPL and AIP, the left LOTc, the pars triangularis of the left inferior frontal gyrus, and the ventral portion of the anterior midcingulate (aMCC) (Fig 2c and Table 2). A cluster within the left ventral premotor cortex also showed a similar trend, though it did not survive the correction for multiple comparisons (k = 244, *p_uncorr_* = 0.045 cluster-level; local maximum at MNI −46, 8, 22; z-score = 3.82; *p_uncorr_* = 0.00006 at voxel-level). As could be anticipated from the literature on spatial and topographic cognition, the comparison of recall of places versus the recall of own paintings revealed greater activation in specific regions including the para-hippocampus, a region recently labeled as the *para-hippocampal place area* (30) (see S5 Fig and S8 Table in the Supporting Information).

**Table 2.**
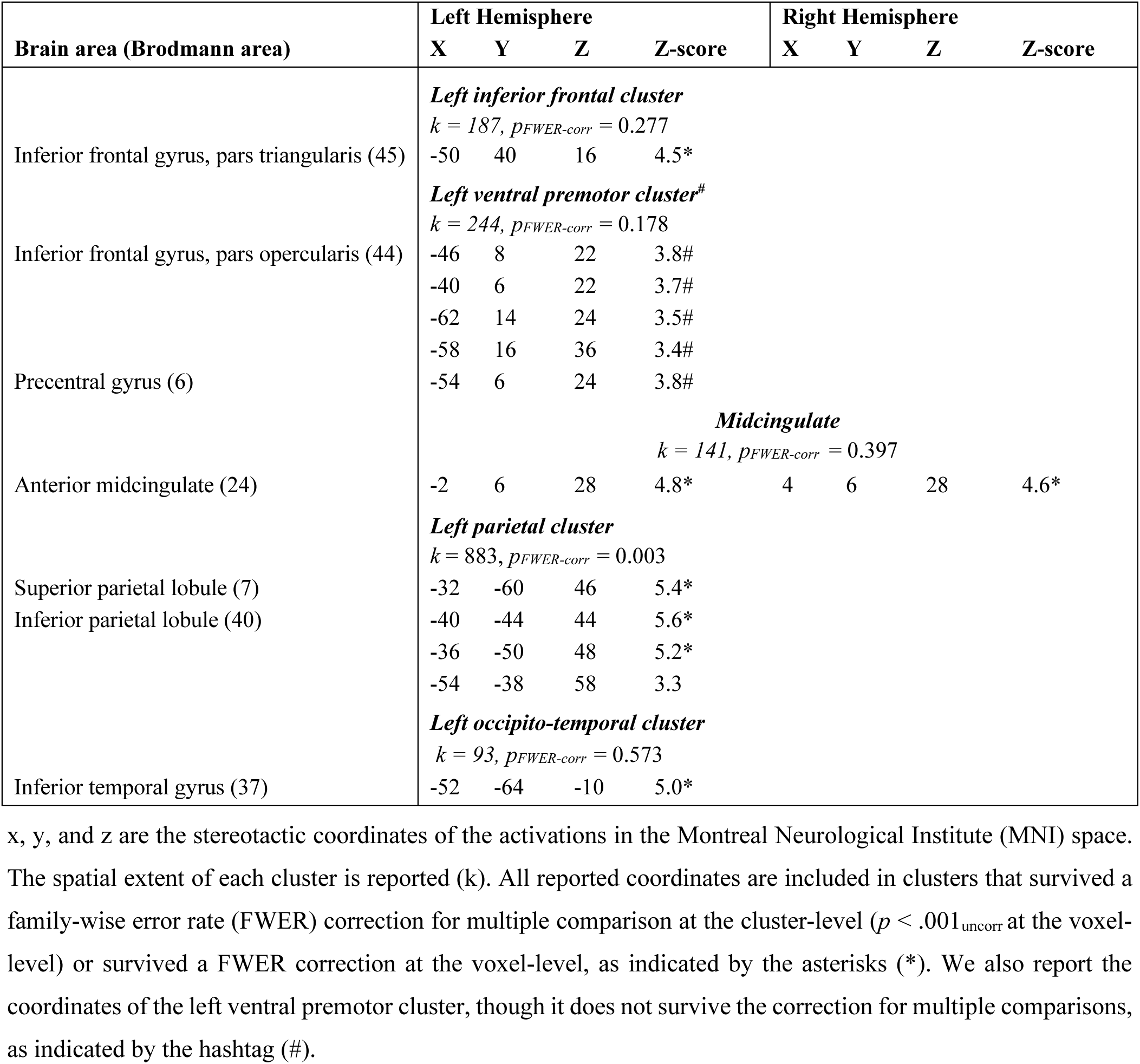
Neurofunctional results of the reconstruction by imagery task (“own paintings > places” masked by the simple effect of own artworks).

Comparing the present findings with the ones of the aesthetic appreciation task in terms of macroscopic distribution of the brain activations, we remark that both reconstructing the creation of one’s own paintings and the appreciation of the same artworks are associated with the recruitment of the left LOTc and the left SPL.

### Neurosynth decoding

We used the Neurosynth.org decoder to qualify further the functional role of the brain regions resulting from the direct comparison between i) the appreciation of one’s own and others’ artworks, and ii) the reconstruction of art creation compared with the control task. Concerning the appreciation of own works of art, besides self-referential anatomical terms like “intraparietal sulcus” (r = 0.303) and “posterior parietal” (r = 0.293), the Neurosynth decoder identified associations with more cognitive-related terms like “attentional” (r = 0.238), “spatial” (r = 0.213), “calculation” (r = 0.191), “working memory” (r = 0.126), “visual” (r = 0.126), and also “**manipulation**” (r = 0.1). Concerning the reconstruction of art creation, besides self-referential anatomical terms like “intraparietal” (r = 0.266), the Neurosynth decoder identified associations with cognitive terms like “calculation” (r = 0.228), “working memory” (r = 0.194), and also “**grasping**” (r = 0.146).

## Discussion

Although at least two alternative views on the neurocognitive bases of art appreciation and production can be found in the literature, no study has so far directly challenged them within the same sample and across both art appreciation and reappraisal of creation. Here, we did so by capitalizing on the opportunity provided by the study of professional painters, who constitute an ideal test-case to explore whether art-making and art-viewing rely on mechanisms of embodied simulations (i.e., *Embodied Hypothesis*) or perceptual learning and inhibitory control (i.e., *Perceptual Learning and Inhibitory Hypothesis*). According to the ***Embodied Hypothesis***, professional artists should show stronger recruitment of brain regions involved in motor resonance **<u>during the observation of their own</u>** versus others’ paintings, as well as while recalling creative moments compared with a well-matched control condition. Instead, according to the ***Perceptual Learning and Inhibitory Hypothesis***, they should show greater recruitment of perceptual and motor inhibitory brain regions **<u>during the observation of others’</u>** than own paintings, as well as the engagement of brain regions involved in inhibitory control during the mental reconstruction of the creative moments.

### Aesthetic appreciation of own and others’ artistic production

Before discussing the results in detail, it is worth recalling that all possible confounds for one’s own versus others’ artworks were carefully controlled for, like the subject – abstract paintings vs. figurative carrying body-related cues – as well as the dynamism or the subjective liking. Accordingly, these factors should not explain the effects discussed below. On the other hand, the self/other distinction was a crucial factor of interest addressed in our study yet entangled with the realm of aesthetic appreciation, rather than based on a generic distinction, to represent how artists make aesthetic judgements on their own work. One could argue that the distinction between self versus other could be confounded with factors like emotional attachment, or autobiographical memory in our artists: however, as far as the aesthetic evaluation was concerned, these hypotheses were made more unlikely by the results of the fMRI investigation (see the discussion below). Having made these preliminary clarifications, we believe that, for aesthetic appreciation, our results are in line with the *Embodied Hypothesis.* The appreciation of one’s own compared with others’ paintings was indeed associated with greater recruitment of brain regions belonging to the so-called action observation network (AON) (31,32) most of which have documented mirror-like properties (32). These regions included the right superior parietal lobule (SPL), with a strong trend in the left-sided region, the right anterior intraparietal sulcus (AIP), the right dorso-central insula, and the left lateral occipitotemporal cortex (LOTc). These areas are systematically recruited in tasks requiring the execution, observation, or imagination of arm/hand actions (31–33) and tool manipulation (34,35). Previous fMRI studies reported the involvement of parietal regions (both AIP and SPL) and LOTc both during the actual manipulation of everyday objects to perform different goal-directed actions (34), as well as during the mental simulation of their use (35). Additionally, a direct neuron-recording study has also provided evidence of the involvement of the posterior parietal cortex (i.e., AIP and SPL) in reaching behaviors (36). The results of the analysis with Neurosynth further support our interpretations. As much as no brain area, with perhaps very few exceptions, is associated with a single function, it is worth emphasizing that the brain map from the aesthetic appreciation task correlated with the term “**manipulation**”. Other correlating terms were “attentional” and “spatial” and “working memory”. Our results also revealed the recruitment of the dorso-central sector of the right insula. Although the insular cortex fulfills different functional roles (i.e., sensorimotor, socio-emotional, olfactory-gustatory, and cognitive), meta-analytic evidence indicates that its dorso-central portion is reliably recruited by sensorimotor tasks (37). This region is anatomically connected with the main nodes of the parieto-frontal circuit involved in reaching/grasping upper limbs movements execution and observation (e.g., with the AIP and the ventral premotor cortex) (38). Interestingly, the dorso-central sector of the right insula has a key role in expressing, perceiving, and imagining the affective component of arm gestures (33). Regardless of their goal, actions can be endowed with different “vitality forms” (33,39). Vitality forms could be defined as the “prosody” of movements, that is, how the gestures are performed (i.e., the style), reflecting the affective state of the agent. The individuals’ internal states modulate the kinematics of actions: the same action can thus be performed in different ways (e.g., gently, vigorously, or neutral) according to the emotional states (33).

These results suggest that, in professional artists, observing their paintings automatically triggers the internal mental representation of the gestures they performed to realize the graphic traces on the canvas. These embodied simulation mechanisms may also extend to the vitality forms and affective states that accompanied and modulated those movements, leaving their imprint in the lines, shapes, and colors of the artwork(40).

Motor simulation processes might also be elicited while appreciating others’ artworks. Indeed, we found the recruitment of bilateral premotor cortices during the observation of both own and other paintings. Yet, embodied mechanisms seem more strongly involved in one’s own works of art. Indeed, own artworks are characterized by a “motor familiarity”, that is, by higher availability and access to the more vivid motor representations of the gestures performed while creating one’s own piece of art. Importantly, these results were independent and therefore not attributable to the different emotional involvement between one’s own and others’ paintings, as confirmed in our control analysis (S4 Table in the Supporting Information). In addition, these observations are also independent of the specific content or stylistic approach of the works of art observed, one’s own or others, as own and others’ artworks were matched for content. The inference that the greater familiarity is visuo-motoric rather than merely pictorial and objectual is supported by the distribution of the activations, which are primarily located in visuo-motor rather than object oriented visual cortices of the ventral stream.

Similar findings are documented by experiments derived from performing arts. Expert dancers show stronger recruitment of mirror areas (e.g., premotor cortex, intraparietal sulcus, and SPL) when they observe dance movements that belong to their motor repertoire compared with kinematically comparable dance movements that do not (41). While expert ballet dancers greatly activate the fronto-parietal mirror system when they see ballet moves compared with capoeira ones, the opposite effect is exhibited by expert capoeira dancers (41), suggesting that these brain responses depend on motor familiarity (i.e., the previous motor experience of performing those actions) and not on visual familiarity (42). In the same vein and more specifically, Calvo-Merino and colleagues (42) performed a fMRI study on male and female classical ballet dancers, showing that the mirror system was more engaged when dancers viewed ballet movements they could represent motorically (i.e., the movements specific to their gender) compared with movements for which they had visual but not motor familiarity (i.e., the ballet movements specific to the other gender, that were visually familiar since female and male dancers train and exhibit together).

Finally, when we explored whether the reward system was modulated by the extent of appreciation of artists’ own and others’ paintings, we identified a cluster within the ventral anterior cingulate cortex and the left nucleus accumbens: these showed differential activations, based on liking judgments, only when artists observed their artworks, with deactivation when artists evaluated own less liked paintings (see Fig 2b). The cluster within the anterior cingulate cortex did overlap with the so-called “field A1” (43–45), a functionally defined region of the medial frontal cortex whose activity correlates with the experience of beauty derived from visual and musical sources (43–45). The nucleus accumbens belongs to the mesolimbic reward circuit and is reliably and directly linked to the experience of pleasure, including in the music domain (46–49). No liking-related modulation was instead present during the observation of others’ artworks.

### Art-making as explored by the reconstruction by imagery task

When participants were asked to mentally reconstruct their past artworks — recalling what appeared in their minds and how they felt during the creative process — a distinct neurofunctional pattern emerged, consistent with the *Embodied Hypothesis*. Brain activations included the left superior parietal lobule (SPL), anterior intraparietal sulcus (AIP), lateral occipitotemporal cortex (LOTc), the pars triangularis of the inferior frontal gyrus (IFG), and the ventral portion of the anterior midcingulate cortex (aMCC). The same trend was also observed in the left ventral premotor cortex.

As previously noted, the SPL, AIP, and LOTc are regions with mirror-like properties involved in executing, observing, and imagining arm and hand movements (32). The pars triangularis of the IFG is likewise engaged during tasks involving the observation or imitation of grasping actions (50,51). Anatomically, this region connects with the SPL and AIP via both the superior longitudinal fasciculus (the dorsal stream) and the extreme capsule fasciculus (the ventral stream) (51).

Finally, the ventral aMCC also plays a central motor role(52). In a large-scale intracortical stimulation study involving over 1,700 sites across the cingulate cortex in 329 patients, Caruana and colleagues provided a comprehensive functional map detailing the specific contributions of each cingulate subregion to behavioral, affective, and sensory functions. (52). Their findings revealed that the stimulation of the aMCC, and particularly its ventral portion, significantly elicited goal-directed behaviors, ranging from simple movements of hands and arms to more complex integrated patterns of reaching and grasping actions performed in a smooth natural way (52). The aMCC also shows mirror-like properties, being involved not only in action execution but also in the observation of actions, especially those expressed through different vitality forms (53). Notably, it is anatomically connected with the dorso-central insula, forming part of a “vitality form” circuit for hand actions (53).

Supporting this interpretation, comparing the brain map generated during art creation imagery with the Neurosynth.org database revealed the association with the term “**grasping**”.

Taken together, our results suggest that motor simulation processes of the kind recruited during motor imagery are involved in artistic production and its recall. In support of this speculation, from the post-scanning debriefing, we learned that more than half of the artists, when reconstructing their moments of creation, visualized themselves in the act of physical creation, rather than in a moment of conceptual conception. Indeed, in addition to remembering the emotions they were experiencing (an aspect common to the re-enactment of the moments when they visited the now familiar places) and the motivations that drove them to realize each artwork or to choose specific colors, twelve out of twenty artists recalled flashes of moments in which they were spreading the color on the canvas, cutting the paper, or sewing fabrics. As some of them indicated, during the art creation process there would not be a sharp distinction between a phase of conception and another of physical implementation, and artworks may arise by implementing motor acts. Coherently, further analyses (see S1 Text in the Supporting Information) confirmed that the regions of the action observation mirror network previously described were recruited by the subgroup of painters who post-scanning declared to employ such motor imagery strategies, while this was not true for the non-motor imagery subgroup (see S6 Fig, S9 and S10 Tables).

The recruitment of sensorimotor regions in visual artists is also consistent with previous results in the musical domain. During musical improvisation, professional musicians show deactivation of prefrontal brain regions involved in executive control (54,55) and a greater involvement of premotor and parietal regions (56). Artistic training could thus lead to lower demands for cognitive control (55,57) and higher recruitment of action simulation processes (56) when creating (and reconstructing the creation of) something new.

### Comparison with previous results and conclusion

While the present results hint that both art-viewing and art-making in professional painters rely on embodied simulation mechanisms, different results (i.e., better aligning with the *Perceptual Learning and Inhibitory Hypothesis*) were found in a recent meta-analysis on the neurofunctional underpinnings of the aesthetic and creative experiences in the visual domain (17). However, several crucial aspects differentiate the present study from previous ones. Only participants naïve to the history or practice of visual art were included in the meta-analysis, and almost all the neuroimaging studies so far applied a “perceptual approach”: artworks were presented for very brief durations (58), and particular care was taken to avoid too easily recognizable pieces of art (17). To bypass the effect of artistic education, experimental subjects were required to evaluate a large number of paintings as beautiful/non-beautiful, independently of previous knowledge. In these instances, the results of the meta-analysis suggested that predictive and inhibitory processes may play an important role in enabling enhanced perceptual processing in “non-expert” participants observing little-known paintings (17). However, expertise influences the emotional and cognitive evaluation of artworks (59,60). Individuals who have been extensively exposed to artistic experiences may have acquired diverse aesthetic sensibility and developed different pathways for appreciating art, which, according to our results, strongly rely also on embodied simulation. Likewise, Alluri et al. (61) demonstrated that, while musicians process the auditory stimuli via an action-based approach during music listening, the non-musicians’ approach is perception-based.

Concerning visual art production, no previous attempt has been made to investigate the neural activities associated with “big creativity”, i.e., artistic production, in professional artists (17), but see (57). Most fMRI studies on visual creativity investigated the neural foundations of what is known as “little creativity” and divergent thinking in naïve participants, asking subjects to find original solutions to open-ended problems under strict time constraints (17,58). Thus, similarly to other domains, different processes and strategies could be employed according to the degree of expertise. While naïve subjects would need to employ inhibitory control over more automatic/common associations, professional artists may be more prone to think out-of-the-box and require such inhibition to a lesser extent. In this view, creative gestures are not the product of artistic creation but can themselves be considered entailed in the creative process, as the artists create-by-doing. As a result, embodied simulation should be seen as the fingerprints of expertise both in visual artistic appreciation and production. In line with the classical dichotomy between the Apollonian (rational) and the Dionysian (corporal) views of art, our results seem to suggest that while artistically naïve individuals may seek conceptual learning while approaching artworks, the corporal dimension is intrinsic to how artists experience their own art. Given the absence of a direct comparison between the data of expert artists and naïve individuals performing similar art-related tasks, this interpretation remains speculative, though we hope it may inspire future research.

The generalizability of our results is subject to some inevitable limitations for the specific population studied: it would be interesting for the future to study aesthetic appreciation in art historians and critics, as they maximally resemble artists for aesthetic expertise while not possessing the pragmatics of an artist. The prediction is made, pending an empirical verification, that they might resemble naïve people than artists while watching works of art, as they should not possess the motoric repertoire needed in art-making. On the other hand, the study of amateur less proficient artists could also be instructive to assess whether the level of proficiency in art making counts in determining patterns of brain activity during aesthetic appreciation.

## Acknowledgments

We are grateful to Fondazione Tecnomed (https://fondazionetecnomed.it/), Paola Rita Lanzoni for her technical support, Massimo Di Domenico, and all the artists who voluntarily took part in this study.

## Supporting Information

**S1 Fig. Trial-timeline of Day1 behavioral tasks.**

**S2 Fig. Behavioral results.**

**S3 Fig. The neurofunctional underpinnings of aesthetic appreciation.**

**S4 Fig. The neurofunctional underpinnings of reconstruction by imagery.**

**S5 Fig. The specific activations for place reconstruction.**

**S6 Fig. The neurophysiology of art experience in the motor imagery subgroup.**

**S1 Table. Neurofunctional results of the simple effect of “own paintings > implicit baseline” during the aesthetic appreciation task.**

**S2 Table. Neurofunctional results of the simple effect of “others’ paintings > implicit baseline” during the aesthetic appreciation task.**

**S3 Table. Neurofunctional results of the conjunction analysis of “own paintings > implicit baseline” ∩ “others’ paintings > implicit baseline” during the aesthetic appreciation task.**

**S4 Table. Neurofunctional results of the simple effect of “own paintings > implicit baseline” during the reconstruction by imagery task.**

**S5 Table. Neurofunctional results of the simple effect of “places > implicit baseline” during the reconstruction by imagery task.**

**S6 Table. Neurofunctional results of the conjunction analysis of “own paintings > implicit baseline” ∩ “places > implicit baseline” during the reconstruction by imagery task.**

**S7 Table. Results of the control analysis related to the neurofunctional differences between own and others’ artworks (“own > others”) irrespective of the differences in emotional involvement.**

**S8 Table. Neurofunctional results of the contrast “places > own paintings” during the reconstruction by imagery task.**

**S9 Table. Neurofunctional results of the aesthetic appreciation task within the subgroup of artists that employed motor imagery strategies during the reconstruction of art creation (“own > others” masked by the simple effect of own artworks).**

**S10 Table. Neurofunctional results of the reconstruction by imagery task within the subgroup of artists that employed motor imagery strategies during the reconstruction of art creation (“own paintings > places” masked by the simple effect of own artworks).**

**S1 Text. Further fMRI analyses on aesthetic appreciation and artistic production.**

