## Supplementary text, tables and figures for "How artists experience their own art"

### Further fMRI analyses on aesthetic appreciation and artistic production

#### The effect of motor imagery strategies

We ran additional analyses to explore whether there were neurofunctional dissociations both on reconstruction of artistic production and the aesthetic appreciation between the sub-group that, during debriefing at the end of the experiment (Day 2), declared to have employed, rather than not, strategies of motor imagery during the reconstruction of art creation. In principle, it is indeed plausible that "motor imagery" artists may exhibit a generally more embodied approach to art than their peers who did not utilize such strategies even when judging works of art. To this end, we replicated the two second-level analyses described in the main text by adding the between-subject factor Group: motor imagery (N =12) vs non-motor imagery (N = 8). This led to two 2x2 ANOVAs having painting (others vs own) as within-subjects factor and group (motor imagery vs non-motor imagery) as between-subjects factor (aesthetic appreciation task), or stimuli (places vs own paintings) as within-subjects factor and group (motor imagery vs non-motor imagery) as between-subjects factor (reconstruction by imagery task). For both analyses we did not find a main effect of group nor an interaction effect. However, we also explored the simple effects "own > others' paintings" (aesthetic appreciation task) and "own paintings > places" (reconstruction by imagery task), separately per each group (motor imagery vs. non-motor imagery) to check whether the results replicated those found on the whole sample. Concerning the aesthetic appreciation task, the simple effects "own paintings > others' paintings" was masked with the simple effect "own paintings > implicit baseline" (voxel-level $p_{uncorr} < 0.05$) calculated within each group. Results revealed that the group of artists that employed motor imagery strategies during the reconstruction of artistic creation showed a greater engagement of the anterior intraparietal sulcus bilaterally during the appreciation of own compared with others' paintings. The same trend was observed within the superior parietal lobule bilaterally (k = 380, $p_{uncorr} = 0.02$ cluster-level; local maximum at MNI -24, -62, 68; z-score = 4.05; $p_{uncorr} = 0.000026$ voxel-level; k = 373, $p_{uncorr} = 0.021$ cluster-level; local maximum at MNI 24, -68, 50; z-score = 3.89; $p_{uncorr} = 0.00005$ voxel-level; Fig. S5a, Table S9). The same contrast did not show any significant results in the other group of artists. Concerning the reconstruction by imagery task, the simple effect "own paintings > places" was masked with the simple effect "own paintings > implicit baseline" (voxel-level $p_{uncorr} < 0.05$) calculated within each group. The results showed that artists who employed motor imagery strategies recruited at a greater extent the lateral occipito-temporal cortex bilaterally, the left superior parietal lobule and intraparietal sulcus, the pars triangularis of the inferior frontal gyrus, and the ventral premotor cortex (Fig. S5b, Table S10). The same contrast did not show any significant results in the other group of artists, possibly due to the small sample size and the different strategies employed among the artists within this group.

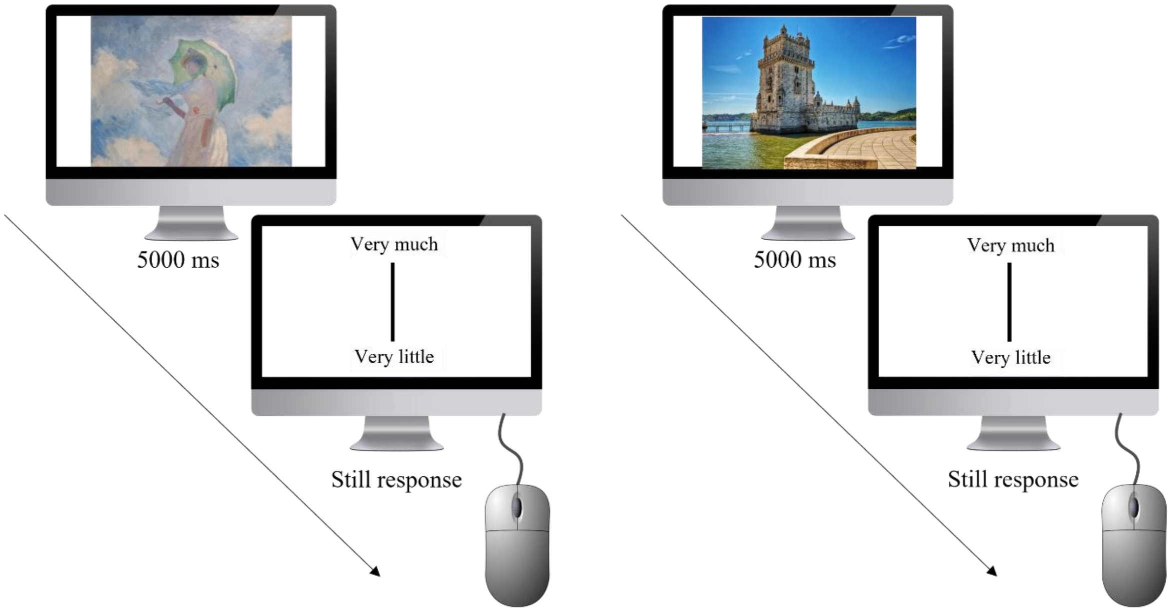


**S1 Fig. Trial-timeline of Day1 behavioral tasks.** The figure illustrates the trial-time line of the two behavioral judgment tasks performed on Day 1. In the first one, artists provided liking judgments on their own and other artists' paintings (left panel). In the second one, they provided liking judgments on places (right panel).

**S1 Table. Neurofunctional results of the simple effect of "own paintings > implicit baseline" during the aesthetic appreciation task.**

| | Left Hemisphere | | | | Right Hemisphere | | | |
|---|---|---|---|---|---|---|---|---|
| **Brain area (Brodmann area)** | **X** | **Y** | **Z** | **Z-score** | **X** | **Y** | **Z** | **Z-score** |
| | *Left superior and medial frontal cluster* *k = 2,058,* $p_{FWER\text{-}corr}$ *< 0.001* | | | | | | | |
| Superior frontal gyrus (8/9) | -16 | 46 | 52 | 4.9* | | | | |
| | -18 | 40 | 56 | 4.8* | | | | |
| | -10 | 60 | 38 | 4.5 | | | | |
| | -10 | 56 | 44 | 4.2 | | | | |
| | -22 | 66 | 16 | 3.3 | | | | |
| Superior medial frontal gyrus (10) | -6 | 64 | 34 | 4.5 | | | | |
| | -8 | 66 | 30 | 4.4 | | | | |
| Supplementary motor area (6) | 0 | 10 | 70 | 4.7* | | | | |
| | -8 | 8 | 74 | 4.7* | | | | |
| Pre-supplementary motor area (6) | -10 | 22 | 66 | 4.3 | | | | |
| | *Left middle-inferior frontal cluster* *k = 2,392,* $p_{FWER\text{-}corr}$ *< 0.001* | | | | *Right middle-inferior frontal cluster* *k = 1,091,* $p_{FWER\text{-}corr}$ *= 0.002* | | | |
| Middle frontal gyrus (9/44) | -56 | 16 | 40 | 5.6* | -- | -- | -- | -- |
| | -42 | 22 | 56 | 3.7 | -- | -- | -- | -- |
| Precentral gyrus (6) | -54 | 10 | 48 | 5.6* | 58 | 8 | 48 | 3.7 |
| | -- | -- | -- | -- | 56 | 6 | 52 | 3.5 |
| Inferior frontal gyrus, pars opercularis (44) | -- | -- | -- | -- | 62 | 16 | 38 | 4.8* |
| | -- | -- | -- | -- | 64 | 16 | 28 | 4.6* |
| | -- | -- | -- | -- | 66 | 14 | 24 | 4.6* |
| Inferior frontal gyrus, pars triangularis (45) | -60 | 28 | 18 | 5.7* | 54 | 40 | 16 | 5.0* |
| | -50 | 42 | 12 | 5.4* | 62 | 30 | 18 | 3.9 |
| | -54 | 32 | 22 | 5.4* | -- | -- | -- | -- |
| | -52 | 42 | 0 | 4.5* | -- | -- | -- | -- |
| Inferior frontal gyrus, pars orbitalis (47) | -36 | 36 | -12 | 4.3 | -- | -- | -- | -- |
| | *Left parieto-occipital cluster* *k = 652,* $p_{FWER\text{-}corr}$ *= 0.015* | | | | | | | |
| Superior parietal lobule (7) | -24 | -64 | 56 | 4.4 | | | | |
| | -24 | -66 | 52 | 4.4 | | | | |
| Middle occipital gyrus (19) | -28 | -64 | 34 | 3.9 | | | | |
| | *Bilateral occipito-temporal cluster* *k = 18,256,* $p_{FWER\text{-}corr}$ *< 0.001* | | | | | | | |
| Superior occipital gyrus (7) | -- | -- | -- | -- | 26 | -62 | 38 | 6.1* |
| Middle occipital gyrus (18) | -28 | -90 | 2 | >8* | -- | -- | -- | -- |
| | -20 | -96 | -2 | >8* | -- | -- | -- | -- |
| | -24 | -96 | 18 | 7.1* | -- | -- | -- | -- |
| Inferior occipital gyrus (18/19) | -30 | -86 | -10 | >8* | 30 | -92 | 0 | >8* |
| | -- | -- | -- | -- | 36 | -78 | -6 | >8* |
| Inferior temporal gyrus (37) | -46 | -60 | -4 | 7.1* | -- | -- | -- | -- |
| Fusiform gyrus (37) | -- | -- | -- | -- | 40 | -60 | -14 | 7.4* |
| | -- | -- | -- | -- | 38 | -44 | -18 | 6.2* |
| Hippocampus (37) | -- | -- | -- | -- | 24 | -28 | -2 | 4.4 |

x, y, and z are the stereotactic coordinates of the activations in the Montreal Neurological Institute (MNI) space. The spatial extent of each cluster is reported (k). All reported coordinates are included in clusters that survived a family-wise error rate (FWER) correction for multiple comparison at the cluster-level ($p < .001_{uncorr}$ at the voxel-level) or survived a FWER correction at the voxel-level, as indicated by the asterisks (*).

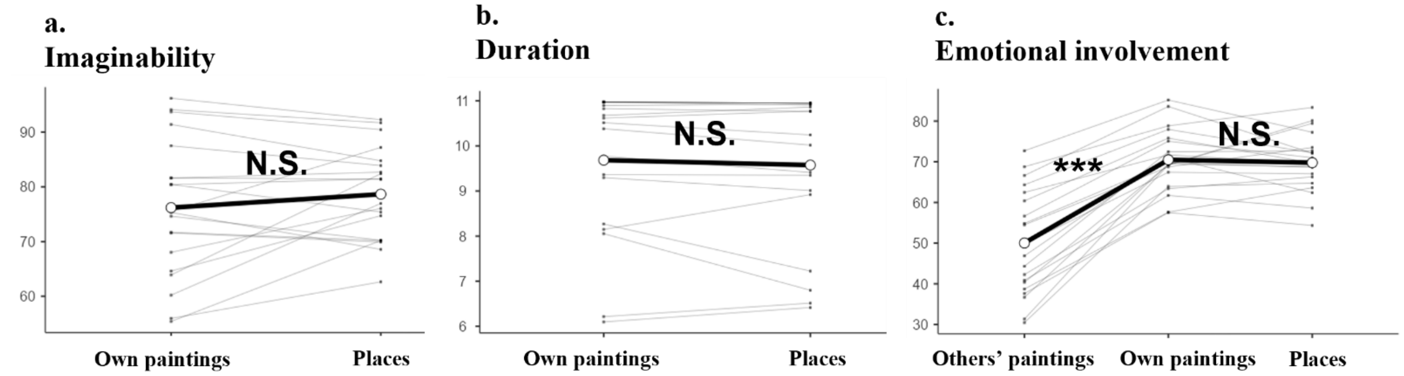


**S2 Fig. Behavioral results.** The figure shows the results of the mixed models analyses on a) imaginability, b) duration, and c) emotional involvement. Random effects are plotted by subjects with grey lines, while fixed effects are indicated with black lines. *** $p < .001$

**S2 Table. Neurofunctional results of the simple effect of "others' paintings > implicit baseline" during the aesthetic appreciation task.**

| Brain area (Brodmann area) | Left Hemisphere X | Y | Z | Z-score | Right Hemisphere X | Y | Z | Z-score |
|---|---|---|---|---|---|---|---|---|
| | *Left superior and medial frontal cluster $k = 1,998$, $p_{FWER\text{-}corr} < 0.001$* | | | | | | | |
| Superior frontal gyrus (8/9) | -12 | 46 | 54 | 5.0* | | | | |
| | -18 | 40 | 56 | 4.7* | | | | |
| | -10 | 62 | 36 | 4.6* | | | | |
| | -18 | 32 | 62 | 4.2 | | | | |
| | -22 | 26 | 66 | 4.1 | | | | |
| Superior medial frontal gyrus (9/10) | -6 | 64 | 34 | 4.7* | | | | |
| | -8 | 60 | 40 | 4.6* | | | | |
| | -8 | 66 | 30 | 4.6* | | | | |
| | -8 | 56 | 46 | 4.5* | | | | |
| Supplementary motor area (6) | -10 | 8 | 78 | 5.4* | | | | |
| | 0 | 10 | 70 | 4.9* | | | | |
| Pre-supplementary motor area (6) | -10 | 22 | 70 | 4.4 | | | | |
| | *Left middle-inferior frontal cluster $k = 2,076$, $p_{FWER\text{-}corr} < 0.001$* | | | | *Right middle-inferior frontal cluster $k = 802$, $p_{FWER\text{-}corr} = 0.007$* | | | |
| Middle frontal gyrus (9/44/45/46) | -56 | 20 | 40 | 5.5* | 52 | 48 | 4 | 4.0 |
| | -50 | 30 | 34 | 4.3 | 50 | 46 | 18 | 3.4 |
| | -46 | 24 | 50 | 4.0 | -- | -- | -- | -- |
| | -42 | 22 | 56 | 3.7 | -- | -- | -- | -- |
| | -38 | 24 | 58 | 3.5 | -- | -- | -- | -- |
| | -46 | 32 | 38 | 3.4 | -- | -- | -- | -- |
| Precentral gyrus (6) | -54 | 8 | 48 | 5.8* | 62 | 14 | 38 | 4.7* |
| | -- | -- | -- | -- | 60 | 14 | 42 | 4.3 |
| | -- | -- | -- | -- | 58 | 8 | 48 | 4.2 |
| Inferior frontal gyrus, pars opercularis (44) | -- | -- | -- | -- | 62 | 20 | 34 | 4.6* |
| | -- | -- | -- | -- | 64 | 18 | 30 | 4.5* |
| | -- | -- | -- | -- | 60 | 20 | 38 | 4.3 |
| | -- | -- | -- | -- | 58 | 24 | 38 | 4.0 |
| Inferior frontal gyrus, pars triangularis (44/45) | -60 | 28 | 18 | 6.3* | 54 | 38 | 20 | 4.6* |
| | -52 | 40 | 14 | 5.4* | 54 | 40 | 16 | 4.6* |
| | -56 | 26 | 28 | 4.9* | 54 | 36 | 24 | 4.5* |
| | -56 | 24 | 32 | 4.9* | 62 | 30 | 18 | 4.3 |
| | -52 | 42 | 0 | 4.7* | 54 | 44 | 6 | 4.0 |
| | -- | -- | -- | -- | 62 | 28 | 22 | 3.9 |
| | -- | -- | -- | -- | 62 | 24 | 20 | 3.8 |
| Inferior frontal gyrus, pars orbitalis (47) | -38 | 34 | -12 | 3.7 | -- | -- | -- | -- |
| | *Bilateral occipito-temporal cluster $k = 18,574$, $p_{FWER\text{-}corr} < 0.001$* | | | | | | | |
| Superior occipital gyrus (7) | -- | -- | -- | -- | 26 | -62 | 38 | 5.2* |
| Middle occipital gyrus (17/18) | -28 | -92 | 2 | >8* | -- | -- | -- | -- |
| | -20 | -96 | -2 | >8* | -- | -- | -- | -- |
| | -14 | -100 | 6 | 7.8* | -- | -- | -- | -- |
| | -24 | -96 | 18 | 7.4* | -- | -- | -- | -- |
| Inferior occipital gyrus (18/19) | -30 | -86 | -10 | >8* | 30 | -92 | 0 | >8* |
| | -- | -- | -- | -- | 36 | -76 | -4 | >8* |
| Inferior temporal gyrus (20) | -50 | -48 | -12 | 4.6* | -- | -- | -- | -- |
| Fusiform gyrus (37) | -38 | -62 | -18 | 6.7* | 38 | -60 | -16 | 7.7* |
| | -38 | -42 | -22 | 5.5* | 38 | -44 | -18 | 6.6* |
| Hippocampus (37) | -- | -- | -- | -- | 24 | -28 | 0 | 4.7* |

x, y, and z are the stereotactic coordinates of the activations in the Montreal Neurological Institute (MNI) space. The spatial extent of each cluster is reported (k). All reported coordinates are included in clusters that survived a family-wise error rate (FWER) correction for multiple comparison at the cluster-level ($p < .001_{uncorr}$ at the voxel-level) or survived a FWER correction at the voxel-level, as indicated by the asterisks (*).

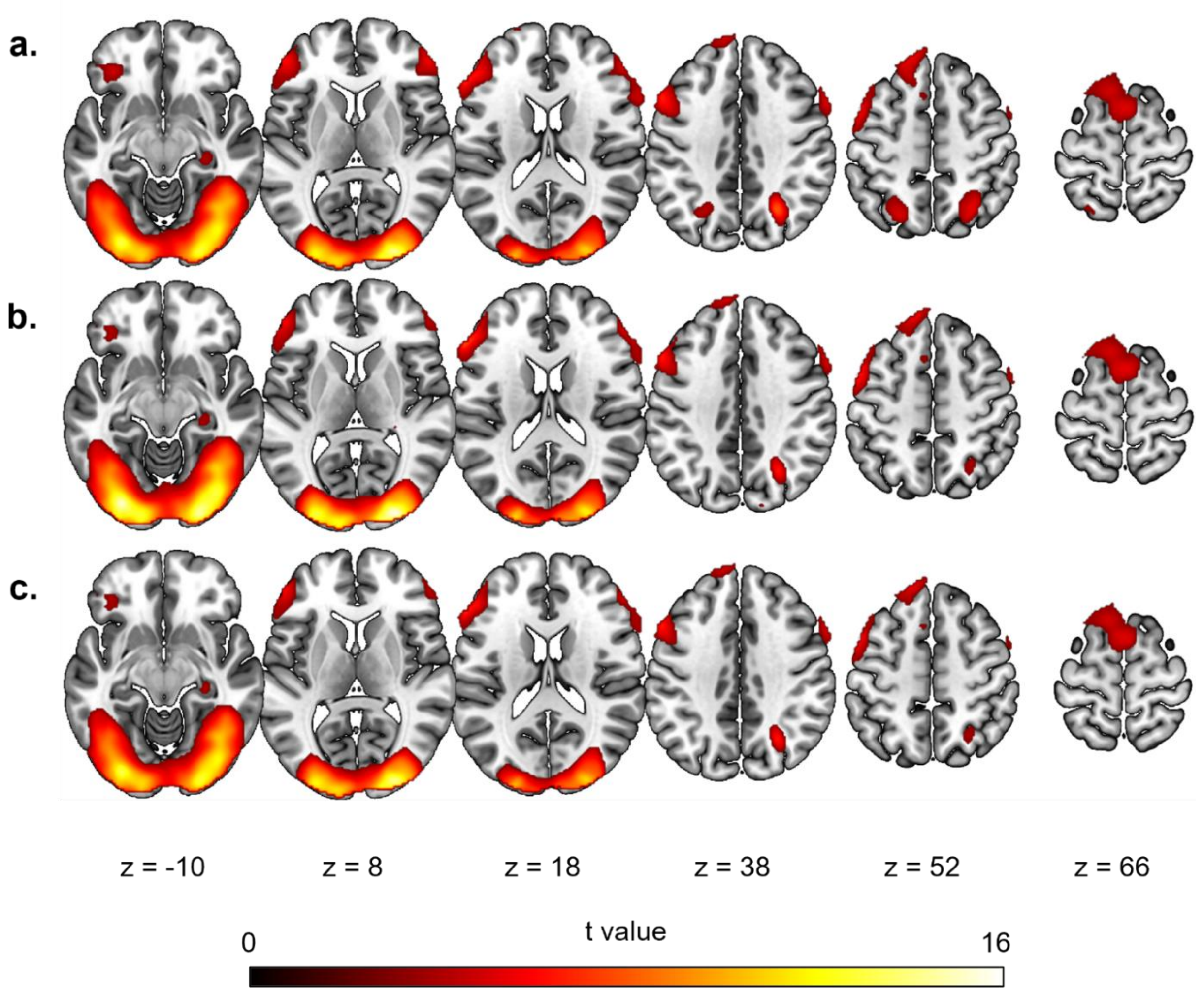


**S3 Fig. The neurofunctional underpinnings of aesthetic appreciation.** The figure illustrates the brain regions involved in a) the simple effect of appreciating own paintings ("own paintings > implicit baseline"); b) the simple effect of appreciating others' paintings ("others' paintings > implicit baseline"); and c) their common activations as measured by a conjunction analysis ("own paintings > implicit baseline" ∩ "others' paintings > implicit baseline"). All the data are reported by applying a statistical threshold of $p_{\text{uncorr}} < .001$ at the voxel level and $p_{\text{FWER-corr}} < 0.05$ at the cluster level.

**S3 Table. Neurofunctional results of the conjunction analysis of "own paintings > implicit baseline" ∩ "others' paintings > implicit baseline" during the aesthetic appreciation task.**

| Brain area (Brodmann area) | Left Hemisphere X | Y | Z | Z-score | Right Hemisphere X | Y | Z | Z-score |
|---|---|---|---|---|---|---|---|---|
| | *Left superior and medial frontal cluster* *k =1,780, $p_{FWER\text{-}corr}$ < 0.001* | | | | | | | |
| Superior frontal gyrus (8/9) | -14 | 46 | 52 | 4.9* | | | | |
| | -18 | 40 | 56 | 4.7* | | | | |
| | -10 | 60 | 38 | 4.5 | | | | |
| | -18 | 32 | 62 | 4.2 | | | | |
| | -10 | 56 | 44 | 4.2 | | | | |
| Superior medial frontal gyrus (10) | -6 | 64 | 34 | 4.5 | | | | |
| | -8 | 66 | 30 | 4.4 | | | | |
| Supplementary motor area (6) | 0 | 10 | 70 | 4.7* | | | | |
| | -8 | 8 | 74 | 4.7* | | | | |
| Pre-supplementary motor area (6) | -10 | 22 | 66 | 4.3 | | | | |
| | *Left middle-inferior frontal cluster* *k = 1,959, $p_{FWER\text{-}corr}$ < 0.001* | | | | *Right middle-inferior frontal cluster* *k = 682, $p_{FWER\text{-}corr}$ = 0.013* | | | |
| Middle frontal gyrus (9/44/45/46) | -56 | 18 | 40 | 5.5* | 52 | 48 | 4 | 4.0 |
| | -42 | 22 | 56 | 3.7 | 50 | 46 | 18 | 3.4 |
| | -38 | 24 | 58 | 3.4 | -- | -- | -- | -- |
| Precentral gyrus (6) | -54 | 10 | 48 | 5.6* | 62 | 14 | 38 | 4.7* |
| | -- | -- | -- | -- | 60 | 14 | 42 | 4.3 |
| | -- | -- | -- | -- | 58 | 8 | 48 | 3.7 |
| | -- | -- | -- | -- | 56 | 6 | 52 | 3.6 |
| Inferior frontal gyrus, pars triangularis (45) | -60 | 28 | 18 | 5.7* | 54 | 40 | 16 | 4.6* |
| | -54 | 32 | 22 | 5.4* | 54 | 44 | 6 | 4.0 |
| | -52 | 40 | 12 | 5.4* | 62 | 30 | 18 | 3.9 |
| | -52 | 42 | 0 | 4.5* | 64 | 24 | 20 | 3.7 |
| Inferior frontal gyrus, pars orbitalis (47) | -38 | 34 | -12 | 3.7 | -- | -- | -- | -- |
| | *Bilateral occipito-temporal cluster* *k = 17,255, $p_{FWER\text{-}corr}$ < 0.001* | | | | | | | |
| Superior occipital gyrus (7) | -- | -- | -- | -- | 26 | -62 | 38 | 5.2* |
| Middle occipital gyrus (18) | -20 | -96 | -2 | >8* | -- | -- | -- | -- |
| | -28 | -90 | 2 | >8* | -- | -- | -- | -- |
| | -24 | -96 | 18 | 7.1* | -- | -- | -- | -- |
| Inferior occipital gyrus (18/19) | -30 | -86 | -10 | >8* | 30 | -92 | 0 | >8* |
| | -- | -- | -- | -- | 36 | -78 | -6 | >8* |
| Inferior temporal gyrus (20) | -50 | -48 | -12 | 4.6* | -- | -- | -- | -- |
| Fusiform gyrus (37) | -- | -- | -- | -- | 40 | -60 | -14 | 7.4* |
| | -- | -- | -- | -- | 38 | -44 | -18 | 6.2* |
| Hippocampus (37) | -- | -- | -- | -- | 24 | -28 | -2 | 4.4 |

x, y, and z are the stereotactic coordinates of the activations in the Montreal Neurological Institute (MNI) space. The spatial extent of each cluster is reported (k). All reported coordinates are included in clusters that survived a family-wise error rate (FWER) correction for multiple comparison at the cluster-level ($p < .001_{uncorr}$ at the voxel-level) or survived a FWER correction at the voxel-level, as indicated by the asterisks (*).

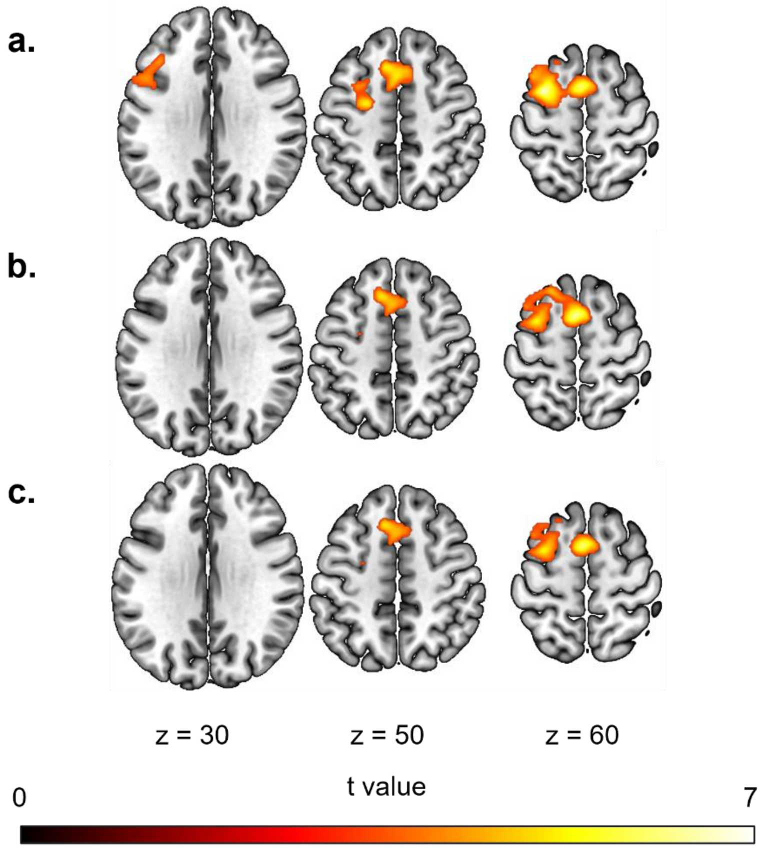


**S4 Fig. The neurofunctional underpinnings of reconstruction by imagery.** The figure illustrates the brain regions involved in: a) the simple effect of recalling the creative moments ("own paintings > implicit baseline"); b) the simple effect of recalling the visiting of known places ("places > implicit baseline"); and c) their common activations as measured by the conjunction effect ("own paintings > implicit baseline" ∩ "places > implicit baseline"). All the data are reported by applying a statistical threshold of $p_{uncorr} < .001$ at the voxel level and $p_{FWER\text{-}corr} < 0.05$ at the cluster level.

**S4 Table. Neurofunctional results of the simple effect of "own paintings > implicit baseline" during the reconstruction by imagery task.**

| Brain area (Brodmann area) | Left Hemisphere X | Y | Z | Z-score | Right Hemisphere X | Y | Z | Z-score |
|---|---|---|---|---|---|---|---|---|
| | *Bilateral frontal cluster* *k = 2,182,* $p_{FWER\text{-}corr}$ < 0.001 | | | | | | | |
| Superior frontal gyrus (6) | -18 | 10 | 70 | 3.8 | -- | -- | -- | -- |
| | -24 | -6 | 52 | 4.7* | | | | |
| Supplementary motor area (6) | -4 | 6 | 58 | 5.2* | -- | -- | -- | -- |
| Posterior medial frontal cortex (8/32) | -8 | 20 | 52 | 4.6* | -- | -- | -- | -- |
| | -4 | 20 | 46 | 4.7* | 8 | 22 | 46 | 4.4 |
| Middle frontal gyrus (6/8) | -30 | 4 | 60 | 5.5* | -- | -- | -- | -- |
| | -34 | 16 | 64 | 4.5 | -- | -- | -- | -- |
| | -30 | 8 | 48 | 3.7 | -- | -- | -- | -- |
| | *Left premotor cluster* *k = 625,* $p_{FWER\text{-}corr}$ *= 0.013* | | | | | | | |
| Precentral gyrus (6) | -54 | 8 | 24 | 3.7 | | | | |
| | -46 | 6 | 44 | 3.6 | | | | |
| Inferior frontal gyrus, pars opercularis (44) | -50 | 16 | 36 | 4.8* | | | | |
| Inferior frontal gyrus, pars triangularis (45) | -38 | 26 | 30 | 3.8 | | | | |
| | | | | | *Right cerebellum* *k = 182,* $p_{FWER\text{-}corr}$ *= 0.28* | | | |
| Cerebellum | | | | | 40 | -72 | -54 | 4.9* |

x, y, and z are the stereotactic coordinates of the activations in the Montreal Neurological Institute (MNI) space. The spatial extent of each cluster is reported (k). All reported coordinates are included in clusters that survived a family-wise error rate (FWER) correction for multiple comparison at the cluster-level ($p < .001_{uncorr}$ at the voxel-level) or survived a FWER correction at the voxel-level, as indicated by the asterisks (*).

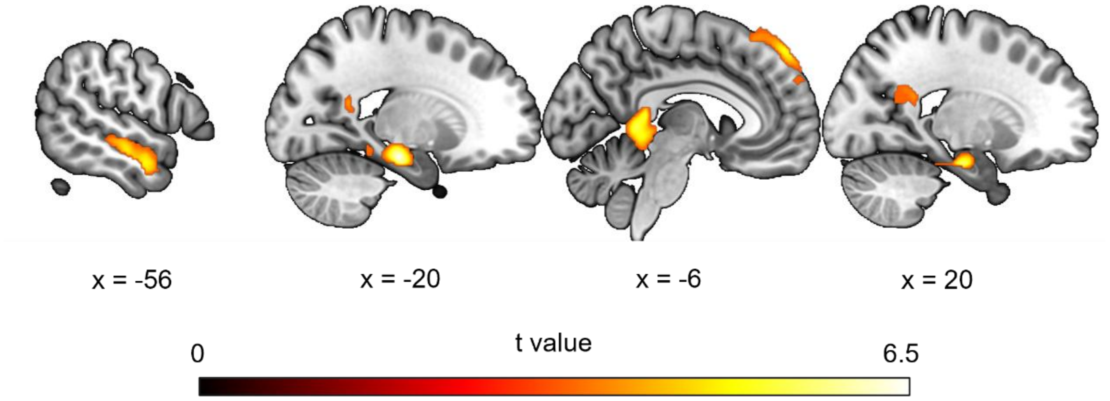


**S5 Fig. The specific activations for place reconstruction.** The figure shows the brain regions that exhibit a greater involvement during the mental reconstruction of the moments when participants visited the places compared with recalling the creative moments (i.e., the effect "places > own paintings"). The statistical threshold was set at $p_{\text{uncorr}} < .001$ at the voxel level and all reported activations survived a family-wise error rate (FWER) correction for multiple comparison at the cluster-level.

**S5 Table. Neurofunctional results of the simple effect of "places > implicit baseline" during the reconstruction by imagery task.**

| Brain area (Brodmann area) | Left Hemisphere | | | | Right Hemisphere | | | |
|---|---|---|---|---|---|---|---|---|
| | X | Y | Z | Z-score | X | Y | Z | Z-score |
| | *Bilateral frontal cluster* $k = 1{,}903$, $p_{FWER\text{-}corr} < 0.001$ | | | | | | | |
| Superior frontal gyrus (6/8) | -20 | 26 | 60 | 4.1 | -- | -- | -- | -- |
| | -18 | 10 | 70 | 3.9 | -- | -- | -- | -- |
| Supplementary motor area (6) | -2 | 8 | 60 | 5.2* | -- | -- | -- | -- |
| | -12 | 8 | 70 | 3.9 | -- | -- | -- | -- |
| Posterior medial frontal cortex (8/32) | -8 | 22 | 52 | 4.7* | -- | -- | -- | -- |
| | -2 | 18 | 48 | 4.3 | 8 | 20 | 48 | 3.9 |
| Middle frontal gyrus (8/9) | -30 | 4 | 60 | 4.6* | -- | -- | -- | -- |
| | -32 | 16 | 66 | 4.0 | -- | -- | -- | -- |
| | -38 | 14 | 62 | 3.8 | -- | -- | -- | -- |
| | -28 | 24 | 60 | 3.5 | -- | -- | -- | -- |

x, y, and z are the stereotactic coordinates of the activations in the Montreal Neurological Institute (MNI) space. The spatial extent of each cluster is reported (k). All reported coordinates are included in clusters that survived a family-wise error rate (FWER) correction for multiple comparison at the cluster-level ($p < .001_{uncorr}$ at the voxel-level) or survived a FWER correction at the voxel-level, as indicated by the asterisks (*).

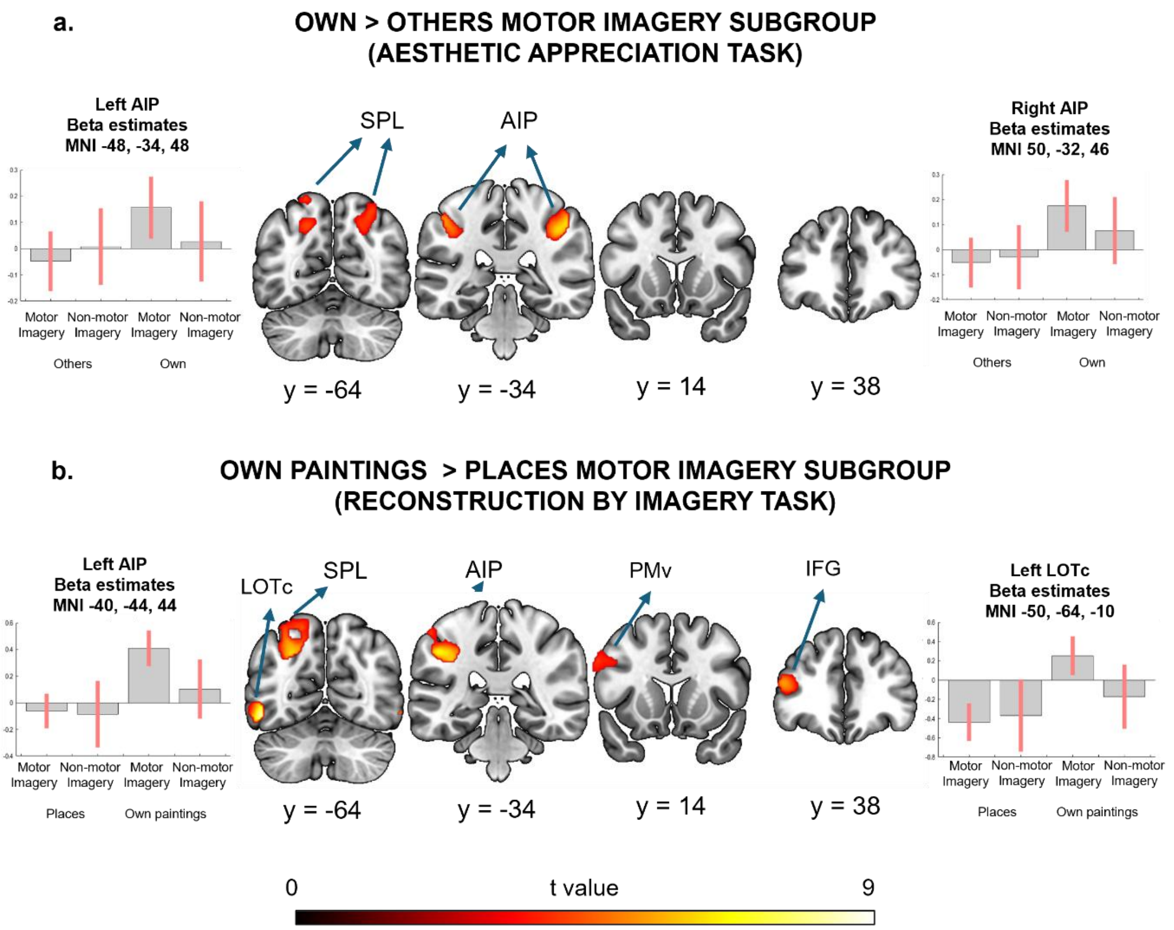


**S6 Fig. The neurophysiology of art experience in the motor imagery subgroup.** The figure illustrates a) the brain regions that show a greater engagement during the aesthetic appreciation of own compared with others' artworks in the motor imagery subgroup, and b) the cerebral regions that exhibit a greater involvement during the reconstruction of the creative moments compared with the control task in the motor imagery subgroup. All the data are reported by applying the same statistical threshold discussed in the text ($p_{\text{uncorr}} < .001$ at the voxel level and $p_{\text{FWER-corr}} < 0.05$ at the cluster or voxel level). For illustrative purposes, we also report the bilateral superior parietal clusters (aesthetic appreciation task) though they do not survive the correction for multiple comparisons. LOTc = lateral occipito-temporal cortex; SPL = superior parietal lobule; AIP = anterior intraparietal sulcus; IFG = inferior frontal gyrus; PMv = ventral premotor cortex.

**S6 Table. Neurofunctional results of the conjunction analysis of "own paintings > implicit baseline" ∩ "places > implicit baseline" during the reconstruction by imagery task.**

| | **Left Hemisphere** | | | | **Right Hemisphere** | | | |
|---|---|---|---|---|---|---|---|---|
| **Brain area (Brodmann area)** | **X** | **Y** | **Z** | **Z-score** | **X** | **Y** | **Z** | **Z-score** |
| | *Bilateral frontal cluster* *k = 1,374,* $p_{FWER\text{-}corr} < 0.001$ | | | | | | | |
| Superior frontal gyrus (6) | -18 | 10 | 70 | 3.8 | -- | -- | -- | -- |
| Supplementary motor area (6) | -2 | 8 | 58 | 5.2* | -- | -- | -- | -- |
| Posterior medial frontal cortex (8/32) | -8 | 20 | 52 | 4.6* | -- | -- | -- | -- |
| | -2 | 18 | 48 | 4.3 | 8 | 20 | 48 | 3.9 |
| Middle frontal gyrus (8/9) | -30 | 4 | 60 | 4.6* | -- | -- | -- | -- |
| | -32 | 16 | 66 | 4.0 | -- | -- | -- | -- |
| | -38 | 14 | 62 | 3.8 | -- | -- | -- | -- |

x, y, and z are the stereotactic coordinates of the activations in the Montreal Neurological Institute (MNI) space. The spatial extent of each cluster is reported (k). All reported coordinates are included in clusters that survived a family-wise error rate (FWER) correction for multiple comparison at the cluster-level ($p < .001_{uncorr}$ at the voxel-level) or survived a FWER correction at the voxel-level, as indicated by the asterisks (*).

**S7 Table. Results of the control analysis related to the neurofunctional differences between own and others' artworks ("own > others") irrespective of the differences in emotional involvement.**

| | Left Hemisphere | | | | Right Hemisphere | | | |
|---|---|---|---|---|---|---|---|---|
| **Brain area (Brodmann area)** | **X** | **Y** | **Z** | **Z-score** | **X** | **Y** | **Z** | **Z-score** |
| | | | | | *Right Insula*# *k = 157,* $p_{FWER\text{-}corr} = 0.317$ | | | |
| Insula | | | | | 40 | -4 | 4 | 3.9# |
| | *Left parietal cluster*# *k = 183,* $p_{FWER\text{-}corr} < 0.250$ | | | | *Right parietal cluster* *k = 1040,* $p_{FWER\text{-}corr} < 0.001$ | | | |
| Superior parietal lobule (7) | -26 | -70 | 48 | 3.3# | 34 | -60 | 54 | 4.1 |
| | -26 | -72 | 56 | 3.3# | -- | -- | -- | -- |
| | -18 | -62 | 44 | 3.3# | -- | -- | -- | -- |
| Precuneus (7) | -14 | -70 | 66 | 3.2# | -- | -- | -- | -- |
| Supramarginal gyrus (40) | -- | -- | -- | -- | 52 | -30 | 44 | 4.2 |
| | *Left occipito-temporal cluster* *k = 273,* $p_{FWER\text{-}corr} = 0.113$ | | | | | | | |
| Inferior temporal gyrus (37) | -50 | -60 | -6 | 5.0* | | | | |

x, y, and z are the stereotactic coordinates of the activations in the Montreal Neurological Institute (MNI) space. The spatial extent of each cluster is reported (k). Coordinates that survived a FWER correction at the voxel-level are indicated by asterisks (*). We also report the coordinates of the right insular cluster and the left parietal cluster, though they do not survive the correction for multiple comparisons ($p < .001_{\text{uncorr}}$ at the voxel-level), as indicated by the hashtag (#).

**S8 Table. Neurofunctional results of the contrast "places > own paintings" during the reconstruction by imagery task.**

| Brain area (Brodmann area) | Left Hemisphere X | Y | Z | Z-score | Right Hemisphere X | Y | Z | Z-score |
|---|---|---|---|---|---|---|---|---|
| | *Bilateral medial frontal cluster* $k = 652$, $p_{FWER\text{-}corr} = 0.011$ | | | | | | | |
| Medial superior frontal gyrus (8/9/10) | -6 | 52 | 52 | 4.5 | 2 | 56 | 32 | 3.4 |
| | -8 | 34 | 60 | 3.6 | 4 | 56 | 36 | 3.4 |
| | -10 | 62 | 30 | 3.3 | -- | -- | -- | -- |
| | -6 | 60 | 32 | 3.2 | -- | -- | -- | -- |
| | *Left temporal cluster* $k = 655$, $p_{FWER\text{-}corr} = 0.011$ | | | | *Right temporal cluster* $k = 229$, $p_{FWER\text{-}corr} = 0.200$ | | | |
| Middle temporal gyrus (21) | -62 | -6 | -16 | 5.1* | -- | -- | -- | -- |
| | -58 | -18 | -12 | 4.3 | -- | -- | -- | -- |
| Parahippocampal gyrus (35) | -- | -- | -- | -- | 20 | -16 | -22 | 4.9* |
| | *Bilateral parieto-occipito-temporal cluster* $k = 2017$, $p_{FWER\text{-}corr} < 0.001$ | | | | | | | |
| Precuneus (17/30) | -16 | -50 | 18 | 3.8 | -- | -- | -- | -- |
| | -20 | -52 | 18 | 3.8 | -- | -- | -- | -- |
| Posterior cingulate cortex (29) | -8 | -40 | 8 | 4.6* | -- | -- | -- | -- |
| Lingual gyrus (27/29) | -8 | -46 | 2 | 4.7* | 8 | -48 | 6 | 4.6* |
| | -- | -- | -- | -- | 10 | -40 | -2 | 4.2 |
| Parahippocampal gyrus (35/37) | -20 | -20 | -18 | 5.3* | -- | -- | -- | -- |
| | -30 | -38 | -10 | 4.1 | -- | -- | -- | -- |
| Cerebellum | -12 | -36 | -14 | 3.4 | -- | -- | -- | -- |

x, y, and z are the stereotactic coordinates of the activations in the Montreal Neurological Institute (MNI) space. The spatial extent of each cluster is reported (k). All reported coordinates are included in clusters that survived a family-wise error rate (FWER) correction for multiple comparison at the cluster-level ($p < .001_{uncorr}$ at the voxel-level) or survived a FWER correction at the voxel-level, as indicated by the asterisks (*).

**S9 Table. Neurofunctional results of the aesthetic appreciation task within the subgroup of artists that employed motor imagery strategies during the reconstruction of art creation ("own > others" masked by the simple effect of own artworks).**

| Brain area (Brodmann area) | Left Hemisphere | | | | Right Hemisphere | | | |
|---|---|---|---|---|---|---|---|---|
| | **X** | **Y** | **Z** | **Z-score** | **X** | **Y** | **Z** | **Z-score** |
| | *Left superior parietal cluster*[#] *k = 380,* $p_{FWER\text{-}corr}$ = 0.077 | | | | *Right superior parietal cluster*[#] *k = 373,* $p_{FWER\text{-}corr}$ = 0.081 | | | |
| Superior parietal lobule (7) | -24 | -62 | 68 | 4.0# | 24 | -68 | 50 | 3.9# |
| | -20 | -62 | 44 | 4.0# | 30 | -64 | 54 | 3.8# |
| | -28 | -60 | 68 | 4.0# | -- | -- | -- | -- |
| | -36 | -58 | 64 | 3.9# | -- | -- | -- | -- |
| | -22 | -72 | 54 | 3.5# | -- | -- | -- | -- |
| | *Left inferior parietal cluster* *k = 235,* $p_{FWER\text{-}corr}$ = 0.209 | | | | *Right inferior parietal cluster* *k = 485,* $p_{FWER\text{-}corr}$ = 0.040 | | | |
| Supramarginal gyrus (2) | -- | -- | -- | -- | 50 | -32 | 46 | 5.5* |
| Inferior parietal lobule (40) | -48 | -34 | 48 | 4.7* | -- | -- | -- | -- |

x, y, and z are the stereotactic coordinates of the activations in the Montreal Neurological Institute (MNI) space. The spatial extent of each cluster is reported (k). Reported coordinates are included in clusters that survived a family-wise error rate (FWER) correction for multiple comparison at the cluster-level ($p < .001_{uncorr}$ at the voxel-level) or survived a FWER correction at the voxel-level, as indicated by the asterisks (*). We also report the coordinates of the two superior parietal clusters, though those did not survive the correction for multiple comparisons, as indicated by the hashtag (#).

**S10 Table**. **Neurofunctional results of the reconstruction by imagery task within the subgroup of artists that employed motor imagery strategies during the reconstruction of art creation ("own paintings > places" masked by the simple effect of own artworks).**

| | Left Hemisphere | | | | Right Hemisphere | | | |
|---|---|---|---|---|---|---|---|---|
| **Brain area (Brodmann area)** | **X** | **Y** | **Z** | **Z-score** | **X** | **Y** | **Z** | **Z-score** |
| | *Left inferior frontal cluster* $k = 286$, $p_{FWER\text{-}corr} = 0.129$ | | | | | | | |
| Inferior frontal gyrus, pars triangularis (45) | -48 | 38 | 16 | 4.7* | | | | |
| | *Left ventral premotor cluster* $k = 558$, $p_{FWER\text{-}corr} = 0.020$ | | | | | | | |
| Inferior frontal gyrus, pars opercularis (44) | -42 | 6 | 24 | 4.6* | | | | |
| Precentral gyrus (44) | -60 | 14 | 32 | 3.7 | | | | |
| | *Left parieto-occipital cluster* $k = 1837$, $p_{FWER\text{-}corr} < 0.001$ | | | | | | | |
| Superior parietal lobule (7) | -22 | -64 | 60 | 4.2 | | | | |
| | -36 | -58 | 64 | 3.3 | | | | |
| Inferior parietal lobule (40) | -40 | -44 | 44 | 6.4* | | | | |
| | -50 | -42 | 56 | 4.5 | | | | |
| Middle occipital gyrus (7) | -26 | -64 | 38 | 5.1* | | | | |
| | *Left occipito-temporal cluster* $k = 489$, $p_{FWER\text{-}corr} = 0.031$ | | | | *Right occipito-temporal cluster* $k = 6$, $p_{FWER\text{-}corr} = 0.964$ | | | |
| Inferior temporal gyrus (37) | -50 | -64 | -10 | 6.3* | 58 | -64 | -8 | 4.8* |

x, y, and z are the stereotactic coordinates of the activations in the Montreal Neurological Institute (MNI) space. The spatial extent of each cluster is reported (k). Reported coordinates are included in clusters that survived a family-wise error rate (FWER) correction for multiple comparison at the cluster-level ($p < .001_{uncorr}$ at the voxel-level) or survived a FWER correction at the voxel-level, as indicated by the asterisks (*).
